# NF-Y controls fidelity of transcription initiation at gene promoters through maintenance of the nucleosome-depleted region

**DOI:** 10.1101/369389

**Authors:** Andrew J Oldfield, Telmo Henriques, Dhirendra Kumar, Adam B. Burkholder, Senthilkumar Cinghu, Damien Paulet, Brian Bennett, Pengyi Yang, Benjamin S. Scruggs, Christopher A. Lavender, Eric Rivals, Karen Adelman, Raja Jothi

## Abstract

Faithful transcription initiation is critical for accurate gene expression, yet the mechanisms underlying specific transcription start site (TSS) selection in mammals remain unclear. Here, we show that the histone-fold domain protein NF-Y, a ubiquitously expressed transcription factor, controls the fidelity of transcription initiation at gene promoters. We report that NF-Y maintains the region upstream of TSSs in a nucleosome-depleted state while simultaneously protecting this accessible region against aberrant and/or ectopic transcription initiation. We find that loss of NF-Y binding in mammalian cells disrupts the promoter chromatin landscape, leading to nucleosomal encroachment over the canonical TSS. Importantly, this chromatin rearrangement is accompanied by upstream relocation of the transcription preinitiation complex and ectopic transcription initiation. Further, this phenomenon generates aberrant extended transcripts that undergo translation, disrupting gene expression profiles. These results establish NF-Y as a central player in TSS selection in metazoans and highlight the deleterious consequences of inaccurate transcription initiation.

## INTRODUCTION

While the sequence, structure, and binding partners of gene promoters have been intensely scrutinized for nearly half a century^1^, how the cell discerns when and where to initiate transcription is still not fully understood^2^. Recent studies have established basic rules regarding spatial arrangements of cis-regulatory elements, ordered recruitment of general transcription factors (GTFs) for transcription pre-initiation complex (PIC) formation, and the role of chromatin in defining the promoter environment^3–8^. One key determinant of active gene promoters is the requirement for an accessible transcription initiation/start site (TSS), characterized by a nucleosome-depleted region (NDR) flanked by two well-positioned nucleosomes (the −1 and +1 nucleosomes)^9^.

Within the NDR, several core promoter elements such as the TATA box and the Initiator (Inr) element exhibit a positional bias relative to the TSS^10,11^ and play important roles in TSS selection. However, the core promoter elements vary from one promoter to the next and can be either absent or present multiple times within a single NDR, suggesting that these elements are not the sole determinants of TSS selection. Thus, how the RNA Polymerase II (Pol II) chooses one transcription initiation site over another remains unclear. In yeast, mutational studies have identified several GTFs and other accessory factors with key roles in TSS selection^12–17^. However, despite greater specificity in TSS selection in metazoans, such accessory factors have yet to be described in higher eukaryotes.

NF-Y, also known as the CCAAT-binding factor CBF, is a highly conserved and ubiquitously expressed heterotrimeric transcription factor (TF) composed of the NF-YA, NF-YB and NF-YC subunits, all three of which are necessary for stable DNA binding of the complex^18–22^. NF-YA, which harbors both DNA-binding and transactivation domains, makes sequence-specific DNA contacts, whereas the histone-fold domain (HFD) containing NF-YB and NF-YC interact with DNA via non-specific HFD-DNA contacts^21,22^. The structure and DNA-binding mode of NF-YB/NF-YC HFDs are similar to those of the core histones H2A/H2B, TATA-binding protein (TBP)-associated factors (TAFs), the TBP/TATA-binding negative cofactor 2 (NC2α/β), and the CHRAC15/CHRAC17 subunits of the nucleosome remodeling complex CHRAC^21^.

NF-Y has an established role in gene regulation through cell type-invariant promoter-proximal binding^23,24^. However, the mechanisms through which NF-Y influences gene expression remain unclear. Several lines of evidence have pointed towards a possible role in the recruitment of chromatin modifiers and/or the PIC to the promoters of its target genes (for a review, see Dolfini et al^19^). Previously, we reported a critical role for NF-Y in facilitating a permissive chromatin conformation at cell type-specific distal enhancers to enable master transcription factor binding^23^. Here, we investigate the role NF-Y plays at gene promoters, what effect it may have on chromatin accessibility and recruitment of the transcription machinery, and how this might impact gene expression.

Through genome-wide studies in mouse embryonic stem cells (ESCs), we find that NF-Y is essential for the maintenance of the NDR at gene promoters. Depletion of the NF-YA protein leads to the accumulation of ectopic nucleosomes over the TSS, reducing promoter accessibility. Interestingly, under these conditions, we find that the PIC can relocate to a previously NF-Y-occupied upstream site, from where it commences ectopic transcription initiation. Remarkably, the resulting ectopic transcript can create novel mRNA isoforms and, in a large number of cases, lead to abnormal translation. Overall, we establish NF-Y’s role in TSS selection and demonstrate its importance in safeguarding the integrity of the NDR at gene promoters.

## RESULTS

### NF-Y promotes chromatin accessibility at gene promoters

Our previous characterization of NF-Y binding sites in ESCs revealed that a majority of these binding sites are located within 500 base pairs (bp) of annotated TSSs of protein-coding genes^23^. Further analysis of our NF-Y ChIP-Seq data revealed a positional bias, with nearly all of the NF-Y binding sites located immediately upstream (median distance 94 bp) of the TSS (**Fig. 1a,b**) and a positive correlation between NF-YA occupancy and CCAAT motif occurrence (**Fig. 1b,c and Supplementary Fig. 1a-c**). To investigate whether NF-Y promotes chromatin accessibility at proximal promoters, as it does at distal enhancers^23^, we used small interfering RNA (siRNA) to knockdown (KD) the DNA-binding subunit *NF-YA* in ESCs (**Supplementary Fig. 1d,e**) and assessed DNase I hypersensitivity at candidate promoters either bound or not bound by NF-Y. Quantitative assessment of the relative “openness” of the probed regions revealed that depletion of NF-YA results in a significant reduction in DNA accessibility at promoters bound by NF-Y but not at promoters without NF-Y binding (**Fig. 1d,e and Supplementary Fig. 1f,g**). In agreement with this, genome-wide assessment of chromatin accessibility using ATAC-Seq confirmed loss in ATAC signal that is specific to promoters targeted by NF-Y (**Fig. 1f,g and Supplementary Fig. 1h,i**). These data suggest that NF-Y helps maintain accessible chromatin at promoters.

**Figure 1.**
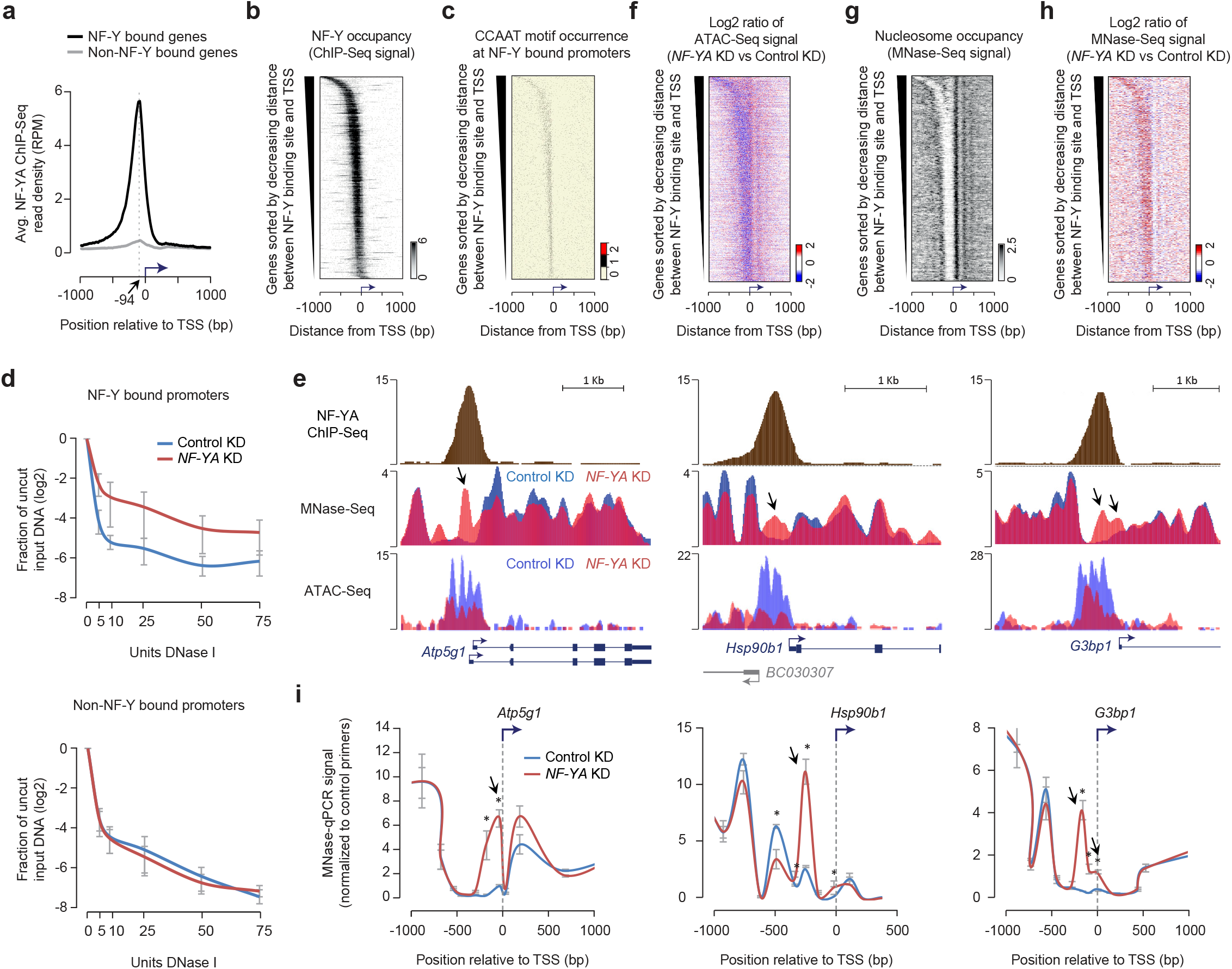
NF-Y binding at gene promoters protects the nucleosome-depleted region (NDR) from nucleosome encroachment. **a,** Average NF-YA occupancy (y-axis), as measured using ChIP-Seq, near transcription start sites (TSSs) of genes with (black, *n*=3,191) or without (grey, *n*=21,195) promoter-proximal NF-Y binding in mouse ESCs. RPM, reads per million mapped reads. **b,** NF-YA occupancy near TSSs of genes with promoter-proximal NF-Y binding. **c,** CCAAT motif occurrence near TSSs of genes with promoter-proximal NF-Y binding. **d,** DNase I hypersensitivity and qPCR analysis of promoters with (*top; n* = 6) or without (*bottom; n* = 4) NF-Y binding in control (blue) and *NF-YA* KD (red) ESCs. Error bars, SEM. Data for individual promoters shown in Supplementary Fig. 1f,g. **e,** Genome browser shots of candidate NF-Y target genes showing nucleosome occupancy in control (blue) or *NF-YA* KD (red) ESCs. Gene structure is shown at the bottom. Arrows highlight nucleosomal gain in *NF-YA* KD cells over what was previously a well-defined NDR in control KD cells. **f,** Relative change (log2) in chromatin accessibility, as measured using ATAC-Seq, near TSSs of genes with promoter-proximal NF-Y binding in *NF-YA* KD vs control KD ESCs. **g,** Nucleosome occupancy (RPKM), as measured using MNase-Seq, near TSSs of genes with promoter-proximal NF-Y binding in control ESCs. RPKM, reads per one kilobase per one million mapped reads. **h,** Relative change (log2) in nucleosome occupancy near TSSs of genes with promoter-proximal NF-Y binding in *NF-YA* KD vs control KD cells. **i,** MNase-qPCR validation of MNase-Seq data shown in Fig. 1f. Error bars, SEM of three biological replicates. *P-value < 0.04 (Student’s t-test, two-sided)

### NF-Y binding protects NDRs from nucleosome encroachment

To further explore the role of NF-Y binding at promoters, we mapped nucleosomes using micrococcal nuclease (MNase) digestion followed by high-throughput sequencing (MNase-Seq). Our data revealed that NF-Y binding sites genome-wide are depleted of nucleosomes (**Fig. 1f and Supplementary Fig. 2a**). Focusing on NF-Y-bound promoter regions, we noted a strong anti-correlation between NF-Y and nucleosome occupancy (**Fig. 1b,h**). To determine whether NF-Y binding plays a direct role in occluding nucleosomes at its binding sites, we performed MNase-Seq in NF-YA depleted cells. Examination of the nucleosomal landscape at candidate NF-Y target promoters in NF-Y depleted cells revealed a striking gain of a nucleosome(s) within what was previously a well-defined NDR (**Fig. 1f,i**), an observation we confirmed by both MNase-qPCR (Fig. 1j) and histone H3 ChIP-qPCR analyses (**Supplementary Fig. 2b**). Genome-wide gain of ectopic nucleosome(s) within the NDR upon *NF-YA* KD is observed specifically at NF-Y bound promoters (**Fig. 1i**), rather than non-NF-Y bound promoters (**Supplementary Fig. 2c,d**), suggesting a direct effect. Interestingly, ectopic nucleosomes observed in NF-YA depleted cells overlap the TSS of NF-Y bound genes (**Fig. 1i and Supplementary Fig. 2d**), suggesting that NF-Y binding could protect the TSS from inhibitory nucleosome binding. Moreover, this ectopic nucleosome positioning in NF-Y depleted cells is consistent with sequence-based predictions of nucleosome binding preference at these regions^25^ (**Supplementary Fig. 2e**). Altogether, these data suggest a role for NF-Y binding at gene promoters in protecting the NDR and the TSS from nucleosome encroachment and that in NF-Y’s absence, nucleosomes are able to bind within this region.

### NF-Y binding impacts PIC positioning and TSS selection

Nucleosomes characteristically provide a refractory chromatin environment for the binding of TFs. In fact, the TATA-binding protein (TBP), and hence the general Pol II transcription machinery, is unable to bind nucleosomal DNA^26^. Consequently, upon observing the appearance of ectopic nucleosome binding within the NDR of NF-Y bound promoters in *NF-YA* depleted cells, we sought to investigate whether this outcome affects binding of the transcription machinery and thus transcription initiation. Because NF-Y is known to interact with the Pol II-recruiting TBP^27^ and several TBP-associated factors (TAFs)^19^, essential components of the TFIID complex that recruits Pol II, we decided to investigate whether TBP enrichment was also affected upon *NF-YA* KD. Indeed, loss of NF-Y binding led to a significant diminishment and/or upstream shift of TBP’s binding pattern (**Supplementary Fig. 3a**), a consequence that could have a dramatic impact on TFIID recruitment, PIC positioning, and thus TSS selection.

In order to obtain a high-resolution view of Pol II activity and to map TSS utilization at base-pair resolution, we performed Start-Seq^9^, a high-throughput sequencing method that captures capped RNA species from their 5’-ends. Start-Seq faithfully mapped the canonical TSSs in the control cells, confirming its utility in capturing transcription initiation sites (**Fig. 2a**). More importantly, consistent with altered TBP binding, analysis of Start-Seq data in NF-YA depleted cells revealed clear upstream shifts in TSS usage at many NF-Y bound promoters (**Fig. 2a**). To ensure identification of promoters exhibiting significant shifts in TSS usage, we used a stringent criterion that excludes ectopic transcription initiation events that occur within ±25bp of the canonical TSS. Our analysis of the 3,056 NF-Y bound gene promoters, using this strategy, identified 538 genes exhibiting significant shifts in TSS location (**Fig. 2b**), with a vast majority exhibiting a TSS shift upstream of the canonical TSS. Ectopic TSSs are located at a median distance of 115bp upstream of the canonical TSS (**Supplementary Fig. 3b**). Importantly, this upstream ectopic transcription initiation is specific to NF-Y bound promoters, with the same directionality of transcription as that from the canonical TSS (**Fig. 2c and Supplementary Fig. 3c**). Interestingly, NF-Y-mediated TSS selection is restricted to TSSs of annotated genes, as the sites of transcription initiation of the associated upstream antisense RNA (divergent transcription^28–30^) or the downstream antisense RNA (convergent transcription) exhibit no such shift (median shift of 0 bp; **Supplementary Fig. 3d**).

**Figure 2.**
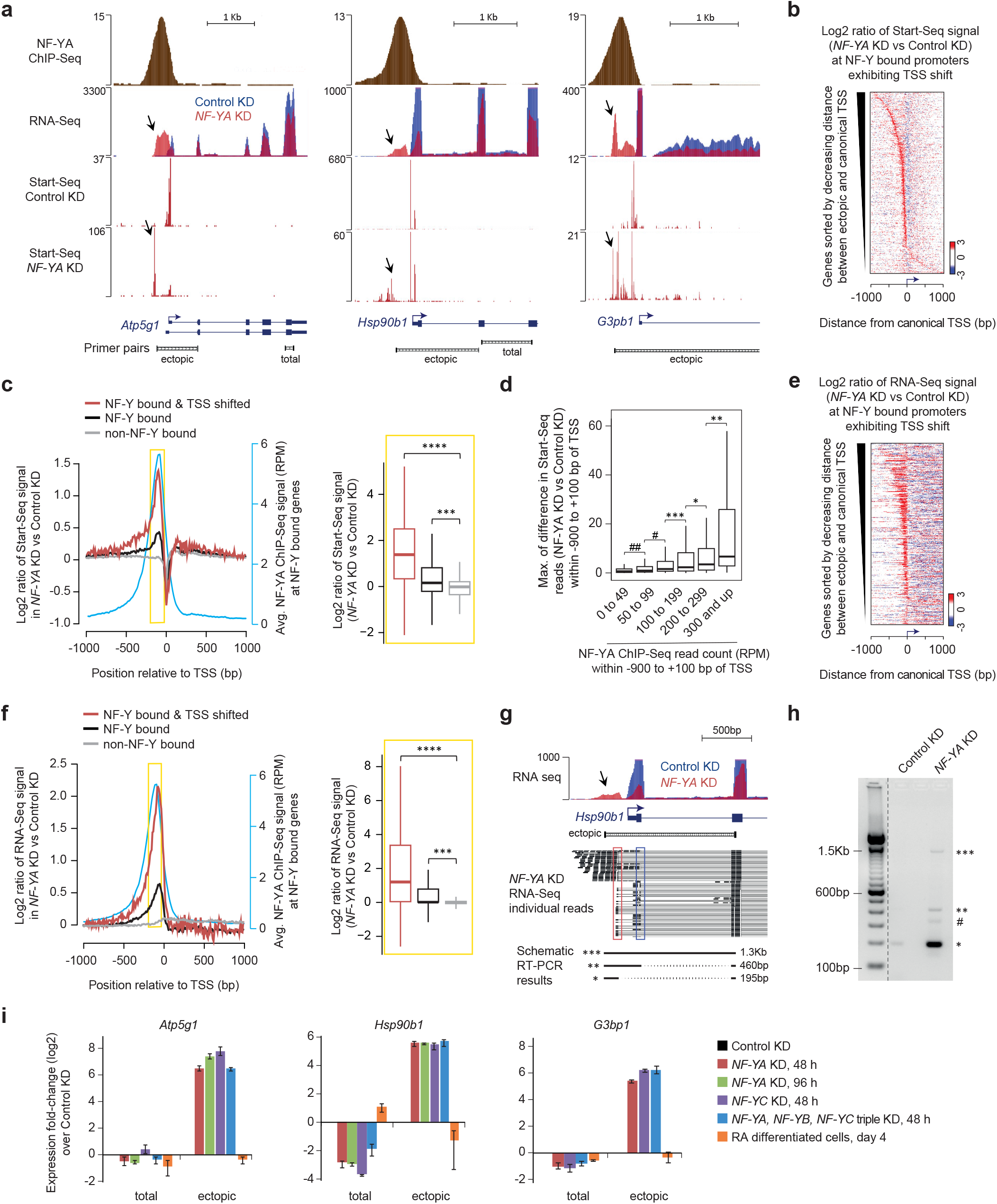
NF-Y binding influences PIC positioning and TSS selection. **a,** Genome browser shots of NF-Y target genes in ESCs showing NF-YA occupancy (ChIP-Seq) and transcription initiation-associated RNA enrichment (Start-Seq) and gene expression (RNA-Seq) in control and *NF-YA* KD ESCs. Arrows highlight regions with ectopic transcription initiation or RNA in *NF-YA* KD cells. Gene structure is shown at the bottom along with PCR amplicons used for RT-qPCR analysis in Fig. 2i. **b,** Relative fold-change (log2) in Start-Seq signal (red, gain; blue, loss) near TSSs of NF-Y bound genes exhibiting TSS shifts (*n* = 538) in *NF-YA* KD vs control KD cells. **c,** *Left:* Average fold-change (log2) in Start-Seq signal near TSSs of NF-Y bound genes exhibiting TSS shifts (red), NF-Y bound genes (black), and non-NF-Y bound genes (gray) in *NF-YA* KD vs control KD cells. Also shown is the average NF-YA occupancy (blue; secondary y-axis) in ESCs. *Right:* Box plot showing the distribution of fold-changes in Start-Seq signal (in *NF-YA* KD vs control KD ESCs) within the upstream proximal-promoter regions (−200 bp to −50 bp; highlighted in yellow). ***P-value = 1.01E-66, ****P-value = 3.53E-128 (Wilcoxon rank-sum test, two-sided) **d,** Box plot showing the distribution of maximum differences in Start-Seq read count between control and *NF-YA* KD cells within the region upstream of TSS (−900 to - 25bp). A 10 bp sliding window was used for computing read count differences. Genes with NF-Y binding were binned into six groups based on NF-YA ChIP-Seq read count within −900 to +100 bp of TSS. *P-value < 0.05, **P-value < 0.0009, ***P-value = 4.47E-07, #P-value = 1.16E-18, ##P-value = 2.49E-18 (Wilcoxon rank-sum test, twosided) **e,** Relative fold-change (log2) in RNA-Seq signal (red, gain; blue, loss) near TSSs of NF-Y bound genes exhibiting TSS shifts (*n* = 538) in *NF-YA* KD vs control KD cells. **f,** *Left:* Average fold-change (log2) in RNA-Seq signal near TSSs of NF-Y bound genes exhibiting TSS shifts (red), NF-Y bound genes (black), and non-NF-Y bound genes (gray) in *NF-YA* KD vs control KD cells. Also shown is the average NF-YA occupancy (blue; secondary y-axis) in ESCs. *Right*: Box plot showing the distribution of fold-changes in RNA-Seq signal (in *NF-YA* KD vs control KD ESCs) within the upstream promoter-proximal regions (−200 to −50 bp; highlighted in yellow). ***P-value = 5.39E-120, ****P-value = 3.44E-158 (Wilcoxon rank-sum test, two-sided) **g, h** Genome browser shot showing RNA-Seq signal at the *Hsp90b1* gene in control (blue) or *NF-YA* KD (red) ESCs (g). Arrow highlights region with ectopic RNA in *NF-YA* KD cells. A representative selection of individual RNA-Seq reads is shown beneath. Red and blue rectangles highlight the ectopic and endogenous splice sites, respectively. Schematic of the RT-PCR results shown at the bottom represent the different PCR amplification products shown in (h). PCR amplification was performed using the “ectopic” primer pair shown underneath the gene structure (g). * denotes new isoform fragment (bypassing the canonical 5’UTR, TSS and the 1st exon), ** denotes mRNA fragment with prolonged 5’ fragment (uses the canonical 1st exon), *** denotes pre-mRNA fragment, and # denotes non-specific/unknown fragment. **i,** RT-qPCR analysis of relative gene expression using the “total” and “ectopic” primer pairs shown in Fig. 2a. Data normalized to *Actin, HAZ* and *TBP*. Error bars, SEM of three to five biological replicates.

Analysis of the differences in Start-Seq read counts upstream of the TSS between control and *NF-YA* KD cells revealed a positive correlation with NF-YA binding intensity (**Fig. 2d**). Comparison of NF-Y-bound promoters that exhibit TSS shifts upon *NF-YA* KD against those that do not revealed stronger NF-YA occupancy at genes with TSS shifts (**Supplementary Fig. 3e**). Moreover, the loss of NF-YA binding at these genes upon *NF-YA* KD results in a substantial increase in nucleosome deposition over canonical TSSs compared to their non-shifting counterparts (**Supplementary Fig. 3f**).

Having established that NF-YA depletion leads to altered localization of the transcription preinitiation complex at a subset of NF-Y-bound promoters, we performed RNA-Seq to assess whether transcription initiation from sites upstream of canonical TSSs gives rise to ectopic transcripts. Consistent with NF-Y having roles beyond that of a canonical transcription activator, loss of NF-YA led to both up- and down-regulation of mRNA levels (**Supplementary Fig. 3g**). Importantly, RNA-Seq experiments in NF-YA depleted cells revealed RNA emanating from sites upstream of the canonical TSS at many NF-Y bound promoters (**Fig. 2e, 2f and Supplementary Fig. 3h**).

Intriguingly, individual paired-end reads from RNA-Seq experiments also revealed that RNAs originating from ectopic TSSs can form stable multi-exonic transcripts (**Fig. 2g**); we confirmed this finding using RT-PCR experiments (**Fig. 2i**). Remarkably, in cases such as *Hsp90b1*, instead of simply extending the 5’ end of the canonical mRNA, the ectopic transcript incorporates a new splicing donor site, resulting in transcripts that skip the canonical TSS, 5’UTR and first exon, giving rise to a new isoform (* product in **Fig. 2h**).

Since hundreds of genes change in expression in response to NF-YA depletion (**Supplementary Fig. 3g**), we wondered if there might be other TFs that could be responsible for the observed effects. To objectively evaluate this possibility, we assessed the enrichment of 680 TF binding motifs (source: JASPAR databse) within the promoter sequences (defined as 200 bp upstream of TSS) of NF-Y bound genes that exhibit a shift in TSS upon NF-YA depletion. Our analysis revealed enrichment of 24 TF binding motifs (**Supplementary Fig. 4a**), of which 14 were also enriched within promoters of non-NF-Y bound genes. Of the 10 TF motifs enriched only within promoter sequences of NF-Y bound genes that exhibit a shift in TSS (**Supplementary Fig. 4b**), eight are not expressed (<1 RPKM) in the control or NF-YA KD ESCs, leaving NF-YA and CREB1 as the only two TFs that could potentially explain the observed effects. The NF-YA motif is present in 88% of the promoters that exhibit a shift in TSS, whereas the CREB1 motif is present in only 33% of the promoters. Furthermore, unlike NF-YA, CREB1 is upregulated (1.6-fold) in NF-YA KD ESCs, making it less likely to explain the TSS shift. Nevertheless, to investigate whether CREB1 motif-containing promoters (that do not bind NF-YA) also exhibit nucleosome encroachment and TSS shift, we examined MNase-Seq and Start-Seq signals in control and NF-YA KD cells and found no obvious changes in the nucleosome positioning or TSS selection (**Supplementary Fig. 4c,d**) to suggest that CREB1 might be responsible for the observed changes at NF-Y bound genes that exhibit a shift in TSS. Collectively, these data support our conclusion that the observed effects are directly attributable to the loss of NF-Y binding.

All our *NF-YA* KD studies were performed 48 hr after siRNA transfection, when NF-YA is depleted but the cells appear normal with no obvious differentiation phenotype^23^. But, to rule out the possibility that the observed changes are due to ESC differentiation as a result of NF-YA depletion, we assessed ectopic TSS usage in ESCs undergoing retinoic acid (RA)-induced differentiation. Four days of RA-induced differentiation did not elicit the use of ectopic TSSs (**Fig. 2i**), indicating that the observed phenomenon is not due to global cellular differentiation effects that may occur upon *NF-YA* KD. Knockdown of *NF-YC*, or all 3 *NF-Y* subunits produced similar results (**Fig. 2i**), making it unlikely that the observed changes are due to potential off-target effects involving siRNA targeting NF-YA. Collectively, these findings suggest that NF-Y binding impacts TSS selection at promoters of protein-coding genes, and that ectopic initiation creates aberrant mRNA species.

### DNA sequence plays an important role in *NF-YA* KD-induced effects

In about 22% of the cases of TSS-shifted genes, such as with the *Ezh2* gene (**Fig. 3a**), the ectopic TSS used in response to *NF-YA* KD corresponds to a previously described alternative TSS, indicating that DNA sequence-based elements play a role in defining sites of ectopic transcription initiation. To further our understanding of how ectopic nucleosome and TSS positioning are established in NF-Y’s absence, we examined the DNA sequence underlying the regions surrounding the canonical and ectopic TSS of NF-Y-bound genes that exhibit TSS shifts upon *NF-YA* KD. *De novo* motif analysis (±5 bp) surrounding the canonical and ectopic TSSs, as determined using Start-Seq, revealed the typical YR initiator dinucleotide^11^ (**Fig. 3b**), indicative of a sequence recognition mechanism determining the location of ectopic TSSs.

**Figure 3.**
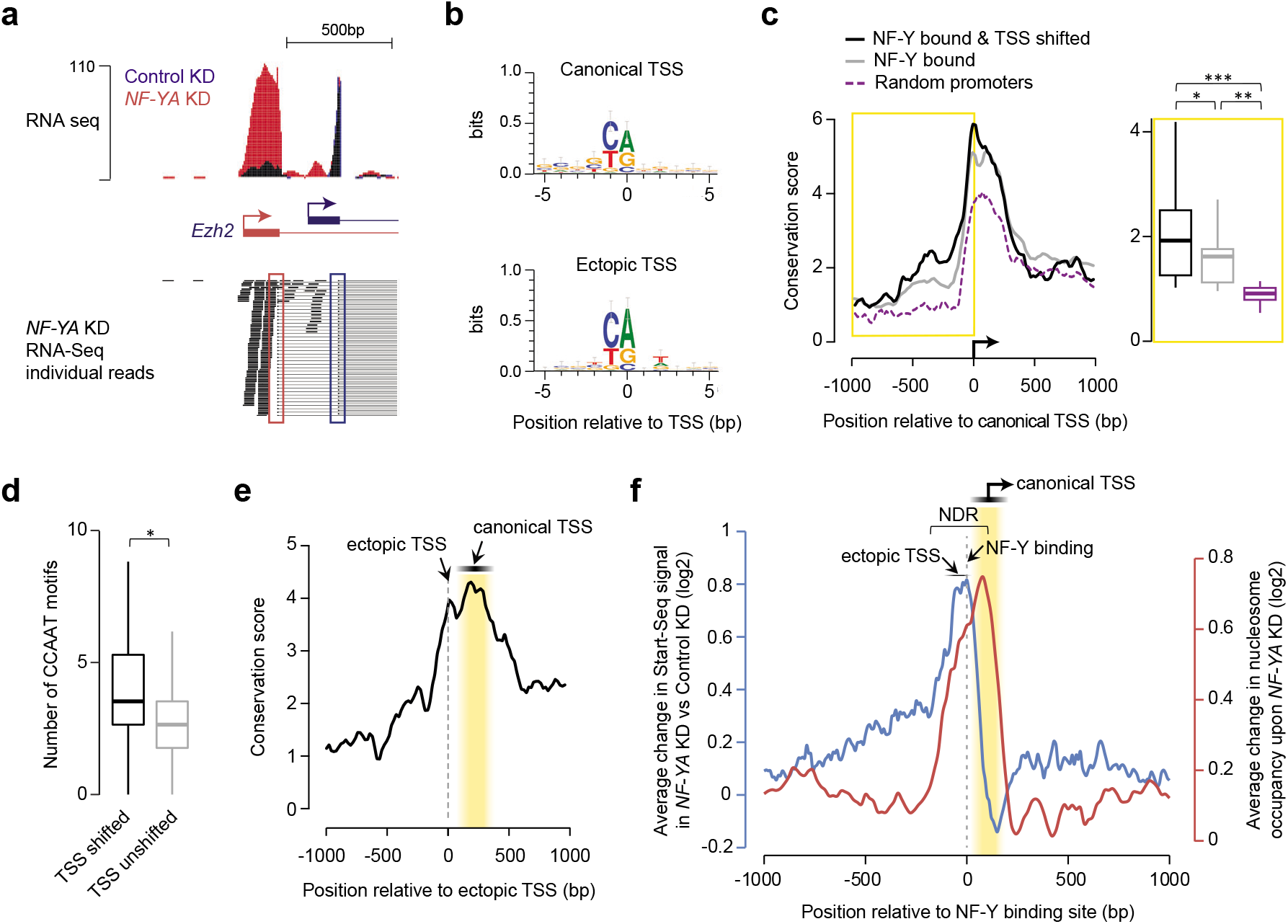
Characterization of ectopic TSSs. **a,** Genome browser shot of RNA-Seq signal at the *Ezh2* gene showing an isoform switch in *NF-YA* KD (red) vs control (blue) ESCs. A representative selection of individual RNA-Seq reads is shown beneath. Red and blue rectangles highlight the ectopic and endogenous splice sites, respectively. **b,** Consensus sequence motif enriched within the 10 bp region centered on canonical TSSs (top) or ectopic TSSs (bottom) of genes with promoter-proximal NF-Y binding, derived from *de novo* motif search. **c,** *Left:* Average sequence conservation in mammals, as computed using PhyloP^74^, of the 2 Kb region centered on canonical TSS of genes with promoter-proximal NF-Y binding (grey, *n* = 3,056), those that bind NF-Y and exhibit TSS shift (black, *n* = 538), or those chosen at random (purple, *n* = 538). *Right:* Box plot showing PhyloP conservation score for the 1 Kb region upstream of canonical TSS. *P-value=0.01372, **P-value=1.69E-10, ***P-value=3.77E-12 (Wilcoxon rank-sum test, two-sided) **d,** Box plot showing the number of CCAAT motifs within 1 Kb region upstream of canonical TSS for genes with promoter-proximal NF-Y binding that exhibit TSS shifts (black) or not (grey) in *NF-YA* KD ESCs. *P-value = 6.67E-36 (Wilcoxon rank-sum test, two-sided) **e,** Average sequence conservation in mammals, as computed using PhyloP^74^, of the 2 Kb region centered on ectopic TSS of genes with promoter-proximal NF-Y binding that exhibit an ectopic TSS (*n* = 538). **f,** Average fold-change (log2; primary y-axis) in Start-Seq signal (blue) in *NF-YA* KD vs control KD ESCs near TSSs of NF-Y bound genes exhibiting TSS shifts (red). Also shown is the average fold-change (log2; secondary y-axis) in nucleosome occupancy (red) in *NF-YA* KD vs control KD ESCs. NDR, nucleosome-depleted region.

We thus we investigated sequence conservation of the regions containing ectopic TSSs. PhyloP conservation scores were calculated from genomic sequence alignment across placental mammals^31^ and NF-Y bound promoters were compared to an equivalent number of randomly chosen gene promoters. This analysis revealed significantly higher conservation of the region immediately upstream of canonical TSSs of NF-Y bound genes (**Fig. 3c**). Further, the conservation at NF-Y bound and TSS-shifted promoters is even higher than that at NF-Y bound promoters in general (**Fig. 3c**), along with a higher enrichment of CCAAT motifs (**Fig. 3d**) and NF-Y occupancy (**Supplementary Fig. 3e**). Focusing our analysis specifically on ectopic TSSs, we discovered a high degree of sequence conservation starting at the ectopic TSS and continuing downstream towards the canonical TSS (**Fig. 3e**), with the NF-Y binding motif more strongly conserved than the regions immediately surrounding it (**Supplementary Fig. 5a**). Therefore, based on the high-level of conservation of NF-Y bound promoter regions, and similarities in NF-Y binding pattern between mouse and human (**Supplementary Fig. 5b**), we propose that NF-Y’s role in the organization of the NDR and in TSS selection is likely conserved in other species.

Altogether, our data, which shows that the ectopic initiation sites are generally located upstream of the NF-Y binding and that the ectopic nucleosomes are observed downstream of the NF-Y binding (**Fig. 3f**), suggest that NF-Y controls the fidelity of transcription initiation at a subset of gene promoters through two complementary mechanisms: (i) NF-Y promotes transcription from the canonical TSS by maintaining the integrity of the NDR, and (ii) NF-Y binding within the NDR per se, either directly or indirectly, prevents PIC from “accidental” utilization of aberrant, upstream sites for transcription initiation.

### RNAs from ectopic TSSs in NF-Y depleted cells undergo translation

Considering the importance of the 5’UTR in the regulation of translation^32^, and the fact that close to 70% of genes showing a TSS shift upon *NF-YA* KD possess an AUG (translation start codon) within the ectopically transcribed region (**Supplementary Table 1**), we explored the potential impact of these aberrant transcripts on translation output. To do so, we performed Ribo-Seq, a ribosome-profiling experiment^33^, on control and NF-YA depleted cells. By sequencing only the ribosome-protected fraction of the transcriptome, Ribo-Seq allows us to determine which RNAs are being actively translated at a given time. Typical of Ribo-Seq, triplet phasing was observed beginning at the annotated translation start site (**Supplementary Fig. 6a**), whereas RNA-Seq presented a flat, uniform distribution (**Supplementary Fig. 6a**). To determine if the transcripts originating from ectopic TSSs in *NF-YA* KD cells are also undergoing translation, we investigated the differences in Ribo-Seq read coverage within the region between the canonical and the ectopic TSSs. At the individual gene level, we can clearly detect the ribosome-protected RNA originating from the region upstream of canonical TSSs of NF-Y bound genes that exhibit ectopic transcription initiation (**Fig. 4a and Supplementary Fig. 6b**). Furthermore, of the 429 NF-Y bound genes with ectopic TSSs that had sufficient Ribo-Seq coverage between the canonical and the ectopic TSSs, 92% showed significantly higher levels of ribosome-protected RNA in *NF-YA* KD cells compared to control KD cells (**Fig. 4b,c**). To ensure that the ribosome-protected RNA, transcribed from the region between the ectopic TSS and canonical TSS, is undergoing translation and is not an artefact, we re-analysed Ribo-Seq data for read coverage phasing after individually determining which ectopically transcribed AUG was most likely to be used as a translation start site for each gene. We found a significant enrichment of triplet periodicity in the Ribo-Seq read coverage beginning at the putative ectopic translation start site, in *NF-YA* KD cells compared to control cells (**Fig. 4d**), indicating that the ectopically transcribed regions indeed undergo translation.

**Figure 4.**
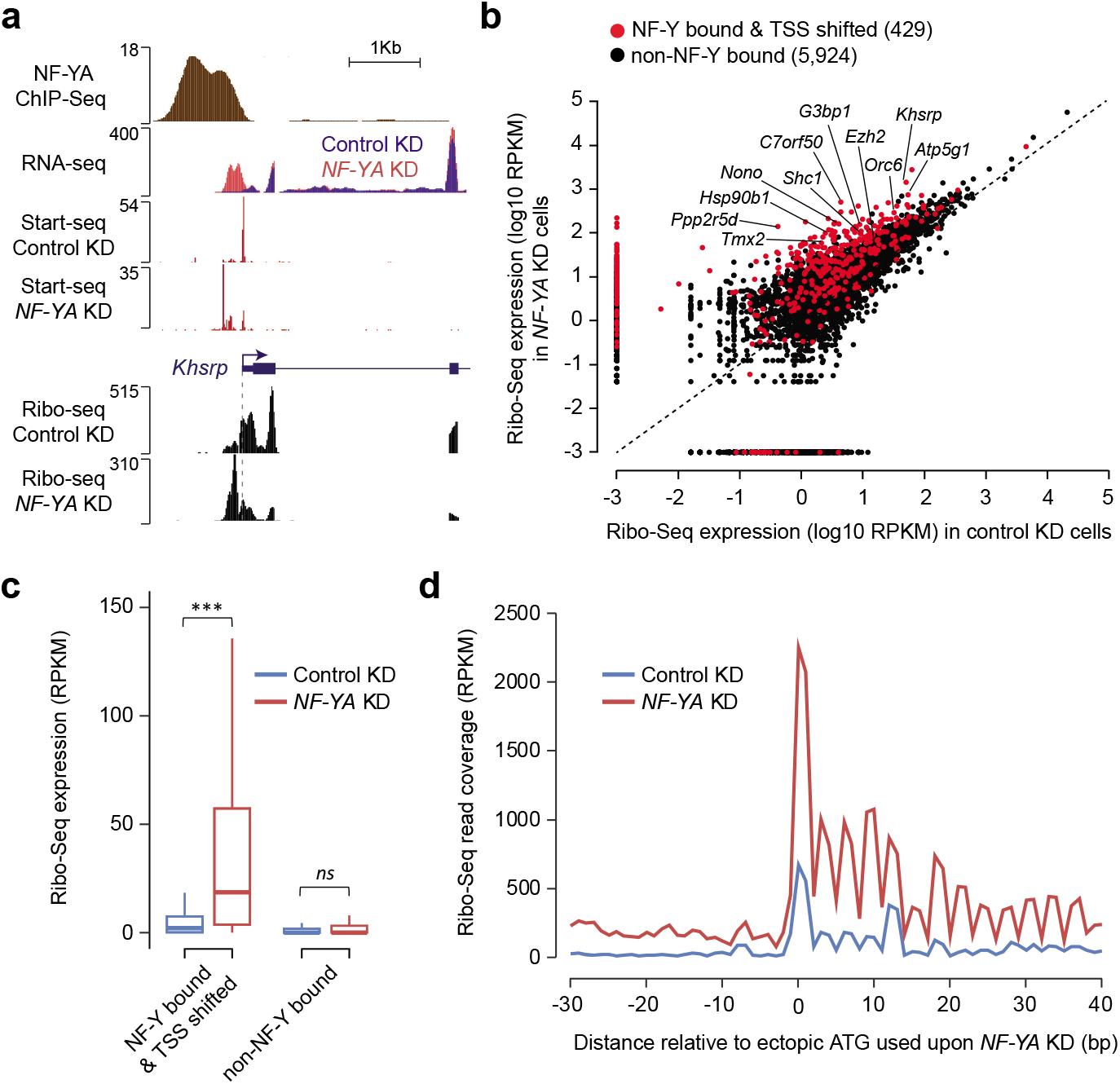
Transcripts originating from ectopic TSSs in NF-Y depleted cells undergo translation. **a,** Genome browser shot of NF-Y target gene *Khsrp* showing ribosome-protected RNA expression, as measured using Ribo-Seq, in control and *NF-YA* KD ESCs. Also shown are tracks for NF-YA ChIP-Seq, and RNA-Seq and Start-Seq in control and *NF-YA* KD ESCs. **b,** Scatter plot showing the Ribo-Seq expression (coverage), in control (x-axis) and *NF-YA* KD (y-axis) ESCs, of the region between the ectopic (shifted) TSS and 25 bp upstream of the canonical TSS of genes with promoter-proximal NF-Y binding that exhibit a TSS shift (red). For comparison purposes, Ribo-Seq expression of the region between 115 bp (median TSS shift distance) and 25 bp upstream of TSS of genes with no promoter-proximal NF-Y binding is shown (black). RPKM, reads per kilobase per million mapped reads. **c,** Box plot showing the Ribo-Seq expression, in control (blue) and *NF-YA* KD (red) ESCs, of the region between the ectopic TSS and 25 bp upstream of the canonical TSS of genes with promoter-proximal NF-Y binding that exhibit an ectopic TSS. For comparison purposes, Ribo-Seq expression of the region between 115 bp (median TSS shift distance) and 25 bp upstream of TSS of genes with no promoter-proximal NF-Y binding is shown. ***P-value = 8.21E-39 (Wilcoxon rank-sum test, two-sided). **d,** Ribo-Seq read coverage, centered on the most likely ectopic start codon (ATG), in control and *NF-YA* KD ESCs. Only genes with promoter-proximal NF-Y binding that exhibit an ectopic TSS in *NF-YA* KD ESCs were used (*n* = 429).

In an attempt to evaluate whether these translated upstream open reading frames (ORFs) generate fusion or variant forms of endogenous protein, we performed western-blot analysis, using commercially available antibodies for candidate proteins, in cells depleted of NF-YA and treated with the proteasome inhibitor MG-132 (to stabilize any unstable fusion proteins; **Supplementary Fig. 6c,d**). We did not detect any obvious aberrant fusion/variant protein of distinct molecular weight. However, in notable cases we did detect altered protein expression levels. For a number of NF-Y bound and TSS-shifted genes, there is a noticeable discordance between the manners with which the RNA and protein levels change in response to *NF-YA* KD (**Supplementary Fig. 6d**). This was surprising considering that changes in transcript levels generally result in proportional changes in protein abundance (Spearman correlation of ~0.8^34, 35^). Recent studies have shown that upstream ORFs (uORFs) can have either a favourable or a deleterious effect on downstream mRNA translation^36–40^ and suggested that the abundance of transcript is less important than the difference in translatability of the canonical versus ectopic transcript^38^. Thus, in situations wherein an upstream extension of the transcript leads to altered protein production, we speculate that this could be due to introduction of an uORF. To summarize, we establish here that transcripts originating from ectopic TSSs undergo translation, and highlight the variable effects this can have on the translation of the canonical ORF within these transcripts.

## DISCUSSION

Driven by the prevalence of the CCAAT motif(s) within core promoters, NF-Y’s function as a regulator of gene expression has almost exclusively been studied in relation to its promoter-proximal binding. Yet, the exact mechanism by which it exerts control over gene expression remains poorly understood. Through comprehensive genome-wide studies in ESCs, we have uncovered a previously unidentified role for NF-Y in safeguarding the integrity of the NDR structure, PIC localization, and TSS selection at protein-coding genes.

### NF-Y binding is incompatible with nucleosome presence

NF-Y can access its target DNA motif, the CCAAT box, in a heterochromatic environment^24^. Furthermore, NF-Y’s unique DNA-binding mode, which induces an ~80° bend in the DNA, may allow and/or promote binding of other TFs, whose recognition sequences become more accessible^23^. Supporting this thesis, DNase experiments have shown that NF-Y is essential for the maintenance of an accessible chromatin^23,41,42^. These attributes have led us and others to propose that NF-Y is a “pioneer factor”^21,23,24,41–43^.

Through comparison of NF-YA ChIP-Seq and MNase-Seq data, we have shown mutual exclusivity between NF-Y and nucleosome occupancy genome-wide (**Fig. 1b,h and Supplementary Fig. 2a**). Given that the structure and DNA-binding mode of NF-YB/NF-YC HFDs are similar to those of the core histones H2A/H2B^21,22,44,45^ our findings suggest steric incompatibility between NF-Y and nucleosomes. This conclusion is supported by the observation that upon *NF-YA* KD, nucleosomes bind within the NDRs left vacant by NF-Y, positioning them in a manner that strongly reflects DNA sequence preferences (**Supplementary Fig. 2e**).

### NF-Y safeguards the integrity of the NDR

The presence of a well-defined NDR within active gene promoters is essential for access by GTFs and PIC assembly, and thus correct transcription initiation. While NF-Y binding does not seem to impact positioning of the +1 or −1 nucleosomes that demarcate the NDR (**Fig. 1f,j,i**), we find that NF-Y is essential for maintaining a nucleosome-depleted NDR. Although we cannot rule out the possibility that NF-Y recruits an ATP-dependent chromatin remodeler to orchestrate nucleosome removal, given NF-Y’s capacity to disrupt the compaction of the chromatin, it is tempting to speculate that NF-Y, with its sequence-specific binding ability, could be acting as an ATP-independent chromatin remodeler. This idea is supported by previous findings that show NF-Y’s capacity to displace nucleosomes in an *in vitro* context^46,47^. Overall, it seems that proteins with tertiary structures similar to core histones can independently preclude nucleosome occupancy. It will be interesting to see if all proteins containing histone-fold domains have a similar effect on nucleosome binding, as has been shown to be the case with subunits of the CHRAC complex^48^.

### NF-Y’s direct and indirect role in PIC positioning

NF-Y has been shown to play a direct role in the recruitment of the pre-initiation complex through interactions with TBP and several TAFs^27,49^. We have shown here that NF-Y also plays an indirect role in the recruitment of PIC-associated proteins, since its binding to promoter-proximal regions is necessary for the maintenance of an open chromatin structure over the TSS, allowing for effective binding of the transcription machinery. In the case of NF-Y’s indirect impact on PIC recruitment, we observed an associated upstream shift in TSS location upon *NF-YA* KD (**Fig. 2b**). Yet, it is interesting to note that only a subset of NF-Y bound genes exhibits this TSS shift. Besides the strength and stability of NF-Y binding and the number of binding events, this likely reflects involvement of additional factors. Our analyses show that efficient utilization of an ectopic TSS requires sequence features within the exposed DNA that are amenable to proper PIC binding, such as previously described alternative start sites (as is the case for *Ezh2*; **Fig. 3c**) or initiator motifs. Moreover, our discovery that a majority of the ectopic TSSs are observed to occur near CCAAT boxes (**Fig. 3c**) suggests that NF-Y binding within the NDR by itself could sterically hinder PIC from aberrant utilization of alternative sites for transcription initiation.

### NF-Y plays an important role in safeguarding transcriptome identity

We thus conclude that NF-Y binding at promoters serves at least three roles: (1) direct PIC recruitment to the promoter region through its interactions with TBP and the TAFs, (2) prevent ectopic nucleosome binding within the NDR through its nucleosome-like structural properties and DNA-binding mode, and (3) occlude alternative transcription initiation sites to ensure correct TSS usage. The shift in TSS usage upon *NF-YA* KD appears to stem from the two latter roles, whereby, upon loss of NF-Y binding, a potential transcription initiation site is uncovered while, simultaneously, a nucleosome prohibits optimal PIC binding to the canonical TSS, forcing the PIC to relocate to an accessible site upstream. Our findings are consistent with studies in transgenic mice showing that the CCAAT-containing Y box sequence is critical for accurate and efficient transcription and that deletion of the Y-box results in aberrant transcripts initiating from regions upstream of canonical TSS^50^.

As might be expected, the usage of an ectopic, upstream TSS has variable consequences on the steady-state levels of resulting mRNAs and protein products. The strength of the ectopic initiation site along with the regulatory potential of the additional upstream mRNA sequences (region between the ectopic and the canonical TSS) undoubtedly impact transcription levels and mRNA stability. Additionally, the TSS employed in NF-YA depleted cells could represent a known alternative start site (as in the case of *Ezh2*; **Fig. 3c**) or generate a previously uncharacterized mRNA isoform (as with *Hsp90b1*; **Fig. 2i**). Furthermore, the upstream extension of mRNA to include a novel ORF can cause abnormal translation (as with *Khsrp* and *C7orf50*; **Fig. 4a and Supplementary Fig. 4b**, respectively). All of these sequelae can affect the quantity and quality of mRNAs and proteins, which could have significant, yet unpredictable, consequences on cell survival or function.

Notably, NF-Y predominantly binds the CG-rich promoters of essential genes (cell-cycle, transcription, DNA repair, etc.), whose accurate expression is vital in most cell types^23,24^. In fact, ~80% of all promoter-proximal NF-Y binding sites overlap CpG islands^23^. Unlike CG-poor promoters, which often correspond to tissue-specific genes and initiate transcription from a well-defined site, CG-rich promoters contain a broad array of transcription initiation sites and often associate with housekeeping genes^2,11,51^. The requirement for NF-Y at promoters of essential genes could reflect both the role of NF-Y in PIC recruitment, and enforcement of appropriate TSS usage.

We suggest that this dual role for NF-Y may explain why promoter-proximal NF-Y binding is so well conserved across mouse and human cell types. Intriguingly, studies in *Saccharomyces pombe* have shown that deletion of *Php5*, an NF-YC orthologue, leads to a ~250bp upstream shift in TSS of the gluconeogenesis gene *Fbp1*^52^. Moreover, in *Saccharomyces cervisiae*, out of the 46 TFs studied, the binding sites for NF-YA homolog Hap2 were shown to have the biggest difference in predicted nucleosome occupancy between Hap2-bound (lower nucleosome occupancy) and non-Hap2 bound (higher occupancy) sites^53^. This opens the exciting possibility of NF-Y’s role in promoter chromatin organization being conserved throughout the eukaryotic kingdom.

### Effects of uORFs within ectopic transcripts on protein expression

A consequence of altered TSS selection upon *NF-YA* KD is that ribosomes can scan over any ectopic mRNA. Ribosomes typically initiate translation upon encountering the AUG start codon, although other codons have been shown to induce translation initiation^54–56^. In our study, we found that nearly three quarters of genes showing a TSS shift upon *NF-YA* KD possess at least one ATG triplet between the ectopic and the canonical TSS. Importantly, we found evidence for translation initiation from such sites (**Fig. 4d**). Given that uORFs can modulate downstream translation and thus act as potent regulators of translation and protein expression^37–40, 57^, it is conceivable that translation initiation from non-canonical start codon(s) within uORFs alters the reading frame and/or protein length; alternatively, it may affect the efficiency with which ribosomes translate the rest of the transcript^58^.

In summary, our studies describe NF-Y’s mechanistic role at promoters, where it is necessary for both maintenance of the NDR’s structural architecture and correct positioning of the transcriptional machinery, therefore influencing TSS selection. Furthermore, our results strongly suggest that the sites of NF-Y binding and the +1 nucleosome demarcate the 5’ and 3’ boundaries, respectively, of the region available for PIC assembly, thereby directing the transcription machinery to the correct TSS while occluding alternative TSSs and other sites of sub-optimal transcription initiation. It will be interesting to explore whether other histone-fold domain proteins, with similar structural and DNA-binding properties analogous to NF-Y, may function in a similar manner.

## METHODS

### Mouse ESC Cell Lines, Culture and RNAi

Mouse ESCs (E14Tg2a; ATCC, CRL-1821) were maintained on gelatin (Sigma, G1890)-coated plates in the ESGRO complete plus clonal grade medium (Millipore), as previously described^23,59^. For experiments, ESCs were cultured on gelatin-coated plates in M15 medium: DMEM (Thermo Fisher, 11965084) supplemented with 15% FBS (Gemini, 100-125), 10 μM 2-mercaptoethanol (Sigma, M3148), 0.1 mM nonessential amino acids (Thermo Fisher, 11140050), 1x EmbryoMax nucleosides (Millipore, ES-008-D), 1 U/ml of ESGRO mLIF (Millipore, ESG1107).

### Transient Transfection

For transfections, ESCs were cultured in M15 medium and transfected with 50 nM siRNA using Lipofectamine 2000 (Thermo Fisher, 11668019) at day 0 and collected after 48 h. Gene-specific siRNAs used: *NF-YA* (Qiagen, SI01327193), *NF-YB* (Invitrogen, MSS247473), *NF-YC* (Qiagen, SI05348217), non-targeting control (Dharmacon, D-001810-02-50).

### Chromatin Immunoprecipitation

ChIP was performed as previously described^23^. Briefly, mouse ESCs (1×10^7^) were cross-linked with 1% formaldehyde (Sigma, F8775) in DMEM (Thermo Fisher, 11965084) for 10 min, and the reaction was quenched by the addition of glycine (Sigma, G8898) at a final concentration of 125 mM for 5 min. Cells were washed twice with PBS, and resuspended in 1 ml of lysis buffer A [50 mM Hepes (Sigma, H3375) pH 7.5; 140 mM NaCl (Sigma, S5150); 1 mM EDTA (Gibco, 15575-038); 10% Glycerol; 0.5% IGEPAL CA-630 (Sigma, I3021); 0.25% Triton X-100 (Sigma, X100); 1x Complete protease inhibitor mixture (Roche, 4693159001), 200 nM PMSF (Sigma, P7626)]. After 10 min on ice, the cells were pelleted and resuspended in 1 ml of lysis buffer B [10 mM Tris-HCl (Sigma, T2663) pH 8.0; 200 mM NaCl (Sigma, S5150); 1 mM EDTA (Gibco, 15575-038); 0.5 mM EGTA (Bioworld, 40520008-2); 1x protease inhibitors (Roche, 4693159001); 200 nM PMSF (Sigma, P7626)]. After 10 min at room temperature, cells were sonicated in lysis buffer C [10 mM Tris-HCl (Sigma, T2663) pH 8.0; 100 mM NaCl (Sigma, S5150); 1 mM EDTA (Gibco, 15575-038); 0.5 mM EGTA (Bioworld, 40520008-2); 0.1% sodium deoxycholate (Sigma, 30970); 0.5% N-lauroylsarcosine (MP, Biomedicals, 190110); 1x protease inhibitors (Roche, 4693159001); 200 nM PMSF (Sigma, P7626)] using Diagenode Bioruptor for 16 cycles (30 sec ON; 50 sec OFF) to obtain ~200–500 bp fragments.

Cell debris were pre-cleared by centrifugation at 14,000 rpm for 20 min, and 25 μg of chromatin was incubated with either NF-YA (Santa Cruz, G-2, sc-17753X), histone H3 (Abcam, ab1791), RNA Pol II (Covance, MMS-126R) or TBP (Abcam, ab51841) antibodies overnight at 4° C. Protein A/G-conjugated magnetic beads (Pierce Biotech, 88846/88847) were added the next day for 2 hours. Subsequent washing and reverse cross-linking were performed as previously described (Heard et al., 2001). ChIP enrichment for a primer-set was evaluated using quantitative PCR, as percentage of input, and normalized to a negative primer-set. See **Supplementary Table 2** for the list of primers used.

### DNase I Hypersensitivity

DNase I hypersensitivity experiments were performed as previously described^23^. Briefly, mouse ESCs treated with non-targeting control siRNA (Dharmacon, D-001810-02-50)or NF-YA siRNA (Qiagen, SI01327193) were collected 48 hours post-transfection in cold PBS. Nuclei were isolated by incubation of 10^7^ cells for 10 min on ice with 5 ml RSB buffer [10 mM Tris–HCl (Sigma, T2663) pH 7.4; 10 mM NaCl (Sigma, S5150); 3 mM MgCl2 (Sigma, M2670); 0.15 mM spermine (Sigma, S3256); 0.5 mM spermidine (Sigma, S0266); 1 mM PMSF (Sigma, P7626); 0.5% IGEPAL CA-630 (Sigma, 13021)], and pelleted by centrifugation at 300 g and 4°C for 10 min. Nuclei were then resuspended in 1 ml DNase reaction buffer [40 mM Tris-HCl (Sigma, T2663) pH 7.4; 10 mM NaCl (Sigma, S5150); 6 mM MgCl2 (Sigma, M2670); 1 mM CaCl2 (Sigma, C1016); 0.15 mM Spermine (Sigma, S3256); 0.5 mM Spermidine (Sigma, S0266)] and counted. Additional resuspension buffer was used to generate equal concentrations of nuclei between samples.

Nuclei from 5×10^5^ cells were aliquoted into microcentrifuge tubes and incubated at 37°C for 5 min with varying amounts of DNase (0 to 75 U; Worthington, LS006344). Digestion was stopped by addition of an equal volume of termination buffer [10 mM Tris (Sigma, T2663) pH 7.4; 50 mM NaCl (Sigma, S5150); 100 mM EDTA (Gibco, 15575-038); 2% SDS (Fisher Scientific, BP166); 10 μg/ml RNAse cocktail (Ambion, AM2286)]. The nuclei were then incubated at 55°C for 15 min, followed by addition of 2 μl of 20 mg/ml Proteinase K (Thermo Fisher, 25530049). Reaction mixtures were incubated overnight at 55°C, followed by a phenol–chloroform extraction of the DNA. The DNA was then precipitated and resuspended in 100 μl H2O. See **Supplementary Table 2** for the list of primers used.

### Quantitative RT-PCR

Quantitative RT-PCR was performed as previously described^23^. Briefly, Total RNAs were prepared from cells using Qiazol lysis reagent (Qiagen, 79306), and cDNAs were generated using the iScript kit (Bio-Rad, 1708891) according to the manufacturer’s instructions. Quantitative PCRs were performed on the Bio-Rad CFX-96 or CFX-384 Real-Time PCR System using the SsoFast EvaGreen supermix (Bio-Rad, 1725201). Three or more biological replicates were performed for each experiment. Data are normalized to *Actin, Haz* and *TBP* expression, and plotted as mean +/- S.E.M. See **Supplementary Table 2** for the list of primers used.

### Western-blot

Western-blots were performed as previously described^23^. Briefly, Cell pellets, lysed in RIPA buffer [25 mM Tris-HCl (Sigma, T2663) pH 7.4; 150 mM NaCl (Sigma, S5150); 1% IGEPAL CA-630 (Sigma, I3021); 1% Sodium deoxycholate (Sigma, 30970)] with protease inhibitors (Roche, 4693159001), were sonicated using Bioruptor (Diagenode) for three cycles (30 sec ON; 50 sec OFF). The lysate was boiled with SDS-PAGE sample buffer (Sigma, S3401), loaded onto NuPAGE gel (Thermo Fisher, NP0321BOX), and transferred using iBlot 2 Transfer Stacks (Thermo Fisher, IB23001). The membranes were then blocked with Odyssey blocking buffer (LI-COR, P/N 927-40000) for 1 h at room temperature with gentle shaking. Each membrane was treated with appropriate primary and secondary (IRDye; LI-COR) antibodies. RNA Pol II and NF-YA levels were measured using an anti-Rpb3 antibody^60^ and anti-NF-YA antibody (G-2; Santa Cruz, sc-17753X) respectively, with Ran (BD Bioscience, 610341) as a loading control. The following antibodies were used in **Figure S4D**: Ppp2r5d (Bethyl, A301-100A), Tmx2 (Abcam, ab105675), Atp5g1 (Abcam, ab180149), Nono (Santa Cruz, sc-166702), C7orf50 (Bethyl, A305-091A), Orc6 (Santa Cruz, sc-390490), Shc1 (Bethyl, A302-019A), Khsrp (Bethyl, A302-021A) and Gapdh (Santa Cruz, sc-25778). The membranes were then washed in PBS (0.1% Tween 20), rinsed with PBS, and scanned and quantified on an Odyssey imaging system.

### MNase-Seq

ES cells were crosslinked for 45 sec with 1% formaldehyde (Sigma, F8775) followed by glycine (125 mM; Sigma, G8898) quenching for 5 min. Cells were washed once with ice-cold PBS and gently resuspended in 300 μl RSB [10 mM Tris HCL (Sigma, T2663) pH 7.4; 10 mM NaCl (Sigma, S5150); 3 mM MgCl2 (Sigma, M2670); 1 mM PMSF (Sigma, P7626); 0.15 mM Spermine (Sigma, S3256); 0.5 mM Spermidine (Sigma, S0266)]. 7ml of RSB + 0.5% IGEPAL CA-630 (Sigma, I3021) were slowly added and the solution was incubated on ice for 10 min. Nuclei were pelleted at 300 X G for 10 min at 4°C and resuspended in 400 μl of digest buffer [15 mM Tris (Sigma, T2663) pH 8.0; 60 mM KCl (Sigma, P9541); 15 mM NaCl (Sigma, S5150); 1 mM CaCl_2_ (Sigma, C1016); 0.25 M sucrose (MP Biomedicals, 152584); 0.5 mM DTT (Roche, 10708984001)], and incubated with 40 U of MNase (Worthington, LS004797) for 5 min. 400 μl stop solution [1% SDS (Fisher Scientific, BP166); 0.1 M sodium bicarbonate (Sigma, S6014); 20 mM EDTA (Gibco, 15575-038)] was added and digested nuclei were incubated at 65°C for 90 min. 24 μl Tris (Sigma, T2663) pH 7.6, 3 μl Proteinase K (Thermo Fisher, 25530049)), and 3 μl GlycoBlue (Ambion, AM9515) were added and digestions were incubated overnight at 65°C. DNA was extracted by phenol–chloroform extraction and mono-nucleosome fragments were gel-purified from fractions digested to ~70% mono-nucleosome, ~20% dinucleosome, and ~10% tri-nucleosome sized fragments. MNase-Seq libraries were prepared using NuGen’s Ovation Ultralow Library System V2 (NuGen, 0344NB-08). See **Supplementary Table 2** for the list of primers used.

### MNase-Seq Data Analysis

MNase-Seq read pairs for all samples were aligned to the mouse (mm9) genome using Bowtie^61^, retaining only uniquely mappable pairs (-m1, -v2, -X10000, --best). Fragments shorter than 120 nt and larger than 180 nt were filtered, as were all duplicate fragments, using custom scripts. Replicates were merged for each condition, and normalized per 10 million uniquely mappable, non-duplicate fragments. BedGraph files containing singlenucleotide resolution fragment centers were generated to facilitate metagene analyses and creation of heatmaps, while whole-fragment coverage bedGraphs were generated for visualization purposes.

### ATAC-Seq

25,000 cells were incubated in CSK buffer (10 mM PIPES pH 6.8, 100 mM NaCl, 300 mM sucrose, 3 mM MgCl2, 0.1% Triton X-100) on ice for 5 min and then centrifuged for 5min. at 4C and 500g. After discarding the supernatant, an aliquot of 2.5 μl of Tn5 Transposase was added to a total 25 μl reaction mixture (TD buffer + H2O). The solution was then heated at 37C for 30min. (with mixing every 10min.). The solution was cleaned up using a MinElute Qiagen kit. After PCR amplification (eight total cycles), DNA fragments were purified with 2 successive rounds of AMPure XP beads (1:3 ratio of sample to beads).

### ATAC-Seq Data Analysis

Low quality reads were removed if they had a mean Phred quality score of <20. Any reads with Nextera adapter sequence were trimmed using cutadapt v1.12. Reads were aligned using Bowtie v1.2 with the following parameters: “-v 2 -m 1 --best --strata”. Reads aligning to the mitochrodia (chrM) were removed, and reads were deduplicated by removing read pairs with both mates aligning to the same location as another read pair. To measure open chromatin, coverage tracks were generated using the first 9 bp of both mates of the aligned reads (corresponding to where the transposase is bound). For smoother coverage tracks that provide better visibility in the genome browser, the original 9 bp regions were extended in both directions an equal distance until the region was 51 bp long. Coverage tracks were normalized to read coverage per 10 million mapped reads (after removing chrM and deduplication).

### RNA-Seq

Total RNA was extracted with Qiazol lysis reagent (Qiagen, 79306) treatment and ethanol precipitation. The samples were then treated with DNAseI Amplification grade (Thermo Scientific, 18068015) and stranded libraries were prepared using the TruSeq stranded RNA kit (Illumina, 20020598) with RiboZero depletion (Gold kit; Illumina, MRZG12324).

### RNA-Seq Data Analysis

Reads were mapped to the mouse (mm9) genome using TopHat v2.1.0^62^. In order to get the transcripts GTF from our samples, Cufflinks^63^ was run with the following options, -g (mm9 GTF from ENSEMBL, version 67, provided as guide). We generated transcriptome assemblies for each of these samples separately and then use Cuffmerge^63^ to combine all the annotations. We used Deseq2^64^ with default parameters for all differential expression analyses with gene count data from Salmon quantification^65^.

### Start-Seq

RNA for Start-Seq experiments were prepared from control or *NF-YA* KD cells as previously described^60^. Briefly, mouse ESCs were grown as described for RNA-Seq and capped-RNAs were isolated essentially as previously described^66^. In brief, approximately 2×10^7^ ESCs were trypsinized and collected by centrifugation. After washing with ice-cold 1x PBS, cells were swelled in 10 ml of Swelling Buffer [10 mM Tris (Sigma, T2663) pH 7.5; 10 mM NaCl (Sigma, S5150); 2 mM MgCl2 (Sigma, M2670); 3 mM CaCl2 (Sigma, C1016); 0.3 M sucrose (MP Biomedicals, 152584); 0.5% IGEPAL CA-630 (Sigma, I3021); 5 mM dithiothreitol (Sigma, D0632); 1 mM PMSF (Sigma, P7626); protease inhibitors (Roche, 4693159001), SUPERase-IN RNAse inhibitor (Ambion, AM2694)] by incubating for 15 min on ice followed by 14 strokes with a loose pestle. The dounced cells were spun for 5 minutes at 500x g, the supernatant (cytoplasm) was discarded, the pellet resuspended in 30 ml of Swelling Buffer and spun as above. The supernatant was discarded and the nuclei pellet was resuspended in 1 ml of Swelling Buffer, aliquoted and stored at −80°C. Libraries were prepared according to the TruSeq Small RNA Kit (Illumina, RS-200-0012). To normalize samples, 15 synthetic capped RNAs were spiked into the Trizol preparation at a specific quantity per 10^6^ cells, as previously described^67^.

### Start-Seq Data Analysis

Start-Seq reads were trimmed for adapter sequence using cutadapt 1.2.1^68^; pairs with either mate trimmed shorter than 20 nt were discarded. A single additional nucleotide was removed from the 3’ end of each read to facilitate mapping of fully overlapping pairs. Remaining pairs were filtered for rRNA and tRNA by aligning to indices containing each using Bowtie 0.12.8 (-v2, -X1000, --best, --un, --max), and retaining unmapped pairs. Following this, a similar alignment was performed to an index containing the sequence of spike-in RNAs only (-m1, -v2, -X1000, --best, --un, --max), and finally, the remaining unmapped reads were aligned to the mouse (mm9) genome utilizing the same parameters, retaining only uniquely mappable pairs.

Strand-specific bedGraph files containing the combined raw counts of short-capped RNA 5’ ends for all control replicates were generated to facilitate observed TSS calling. For all other purposes, 5’ end counts were normalized per 10 million mappable reads, then based on depth-normalized counts aligning to spike-in RNAs. Spike normalization factors were determined as the slope of the linear regression of each sample’s depth normalized spike-in read counts versus the single sample with the lowest total count. Control and NF-YA knockdown bedGraph files were generated from these spike-normalized counts by taking the mean of all replicates, genome-wide, at single-nucleotide resolution.

### Observed TSS Calling

Observed TSSs were identified as previously described^9^, based on the control Start-Seq data, using mm9 RefSeq annotations downloaded from the UCSC genome browser (January 2015). Briefly, the position with the highest read count within 1000 nt of an annotated TSS, or that with the highest count within the 200 nt window of highest read density was selected, depending on proximity. When insufficient Pol II ChIP-Seq signal existed in the 501 nt window centered on the selected locus, relative to a comparable window about the annotated TSS (a ratio less than 2:3), the observed TSS was shifted to the location with the highest Start-Seq read count within 250 nt of the annotated TSS. When fewer than 5 reads were mapped to the selected locus, the annotated TSS was maintained. Groups of transcripts with identical observed TSS were filtered, maintaining a single representative with the shortest annotated to observed TSS distance. Groups of observed TSSs within 200 nt of one another were similarly reduced by first removing any RIKEN cDNAs, predicted genes, or observed TSSs moved to the annotation due to lack of Start-Seq reads. Following this, a single observed TSS was selected based on observed to annotated proximity. In this manner, observed TSSs were called for 24,498 transcripts; of these, 16,483 were selected based on Start-Seq data, while for 8,015 the annotated position was maintained. NFY-bound promoters were then identified as those with an NF-YA ChIP-Seq peak intersecting the observed TSS −900 to +100 nt window.

### Ectopic TSS Calling

Ectopic TSSs were identified through the comparison of NF-YA knock-down Start-Seq read counts to control using DESeq^69^. Counts were determined for all samples in 10 nt bins tiling the region −995 to +995 nt, relative to each observed TSS. Bins closer to an upstream or downstream TSS than their own were excluded, as were those in the observed TSS −25 to +24 nt region. Normalization was performed based on size factors calculated within DESeq from spike-in RNAs alone, ensuring these values were equivalent across all samples. All bins with a positive log2 fold change and adjusted p-value less than 0.1 were identified. If more than one bin associated with a single observed TSS was selected, that with the lowest adjusted p-value was retained. Within each of these bins, the position with the total Start-Seq read count across all NF-YA knock-down samples was selected as the ectopic TSS. In cases where multiple sites exist with identical counts, that closest to the observed TSS was selected.

### Ribo-Seq

Approximately 8-9 × 10^6^ ESCs (per sample) were treated with cycloheximide (0.1 mg/ml; Sigma, 01810-1G) for 1 minute prior to trypsinization and cell lysis. Control and *NF-YA* KD cells were used as input material for the TruSeq Ribo Profile Mammalian Library Prep Kit (Illumina) following the manufacture’s protocol.

### Ribo-Seq Data Analysis

Total RNA-Seq and Ribosome-protected-RNA-Seq (Ribo-Seq) read pairs were trimmed with cutadapt^68^; fragments shorter than 15 nt were discarded. Read pairs were filtered for rRNA and tRNA by aligning to respective indices using Bowtie 0.12.8 (-v2, -X1000, --best, --un, --max), and retaining unmapped pairs. The remaining read pairs were aligned to the mouse (mm9) genome using STAR v2.6.0c^70^. The read counts intersecting CDSs were determined per sample. We then determined the normalization factors from the aligned counts using DESeq2 v1.18.1^64^. Using STAR’s bedGraph output (read pairs with unique alignment), both sequencing runs were merged using unionBedGraphs (a component of bedtools 2.25.0). We then applied the previously calculated normalization factors to the merged bedGraphs. To obtain bigwig format files, both normalized biological replicates were merged using unionBedGraphs and then converted to bigWig. To infer position of ribosomes from Ribo-Seq reads, we used the 5’ end of each fragment. We then added 12 nt in order to locate the highest peak on the A of ATG. The same process was applied to reads from total RNA-Seq for comparison, to check that RNA-Seq read positions do not exhibit a 3n periodicity like Ribo-Seq reads do. To study the coverage in shifted regions we used a two-step process. First, we searched for all the ATGs in those regions. Then, the ATG with the highest coverage within the downstream 60 nt was selected. We finally plotted the cumulative coverage, taking the A of the selected ATG as the anchor. Genes whose cumulative Ribo-seq signal, within the shifted region, represented > 5% of total Ribo-Seq reads were removed from the phasing analysis (*n* = 3).

### Motif Analysis

Around observed TSSs bound by NF-YA, those not bound by NF-YA, those bound by NF-YA with an associated ectopic TSS, as well as bound and non-bound ectopic TSSs themselves, *de novo* motif discovery was performed in the TSS +/- 50 region using MEME^71^ (-dna -mod zoops -nmotifs 25 -minw 6 -maxw 20 -revcomp). TRAP^72^ was used to search for the enrichment of 680 known TF motifs, obtained from the JASPAR database^73^, within the promoter sequences (defined as 200 nucleotides upstream of the TSS). Statistical significance for enrichment of sequence motifs within promoters of interest were calculated in reference to promoter regions from all mouse genes. Benjamini-Hochberg method was used for multiple-testing correction.

### Sequence Conservation and Predicted Nucleosome Occupancy Data

Per-nucleotide predicted nucleosome occupancy for the mouse (mm9) genome was obtained from the authors of a previously published study^25^. Per-nucleotide phyloP conservation scores, based on 20 placental mammals, were downloaded from the UCSC Genome Browser. Both data sets were converted to bedGraph format using custom scripts to facilitate generation of metagene analyses and heatmaps.

### Data Availability

ATAC-Seq, MNase-Seq, RNA-Seq, Start-Seq, and Ribo-Seq data generated for this study have been deposited in the GEO repository under the accession number <u>GSE115110</u>. The NF-YA ChIP-Seq data used in this study, generated for our previous study^23^, can be obtained from the GEO repository under the accession number <u>GSE56838</u>.

## ACKNOWLEDGEMENTS

We thank R. Luco, E. Nora, and P. Navarro for critical comments on the manuscript. We thank A.E. Conway and D. Fargo for useful discussion, J.P. Villemin and E. Agirre for guidance on bioinformatic analyses and F. Aubignat for assistance with figure illustration. We thank the NIEHS Epigenomics core, especially J. Malphurs, for their support. D.P. and E.R. thank the ATGC bioinformatics platform in Montpellier. D.P. and E.R. is supported by the Institut de Biologie Computationnelle (ANR-11-BINF-0002) Institut Français de Bioinformatique (ANR-11-INBS-0013), GEM Flagship project funded from Labex NUMEV (ANR-10-LABX-0020), and France Génomique. This work was supported by the Intramural Research Program of the NIH, National Institutes of Environmental Health Sciences (R.J.: 1Z1AES102625, K.A.: 1ZIAES101987).

## AUTHOR CONTRIBUTIONS

A.J.O., K.A., and R.J. conceived the study. A.J.O., T.H., and S.C. performed the experiments. A.J.O., D.K., A.B.B., D.P., P.Y., B.S.S., C.A.L., B.B., and E.R. analyzed the data. A.J.O., T.H., K.A., and R.J. wrote the manuscript. All authors reviewed, edited, and approved the final version of the manuscript.

## SUPPLEMENTARY INFORMATION

**Supplementary Figure 1.**
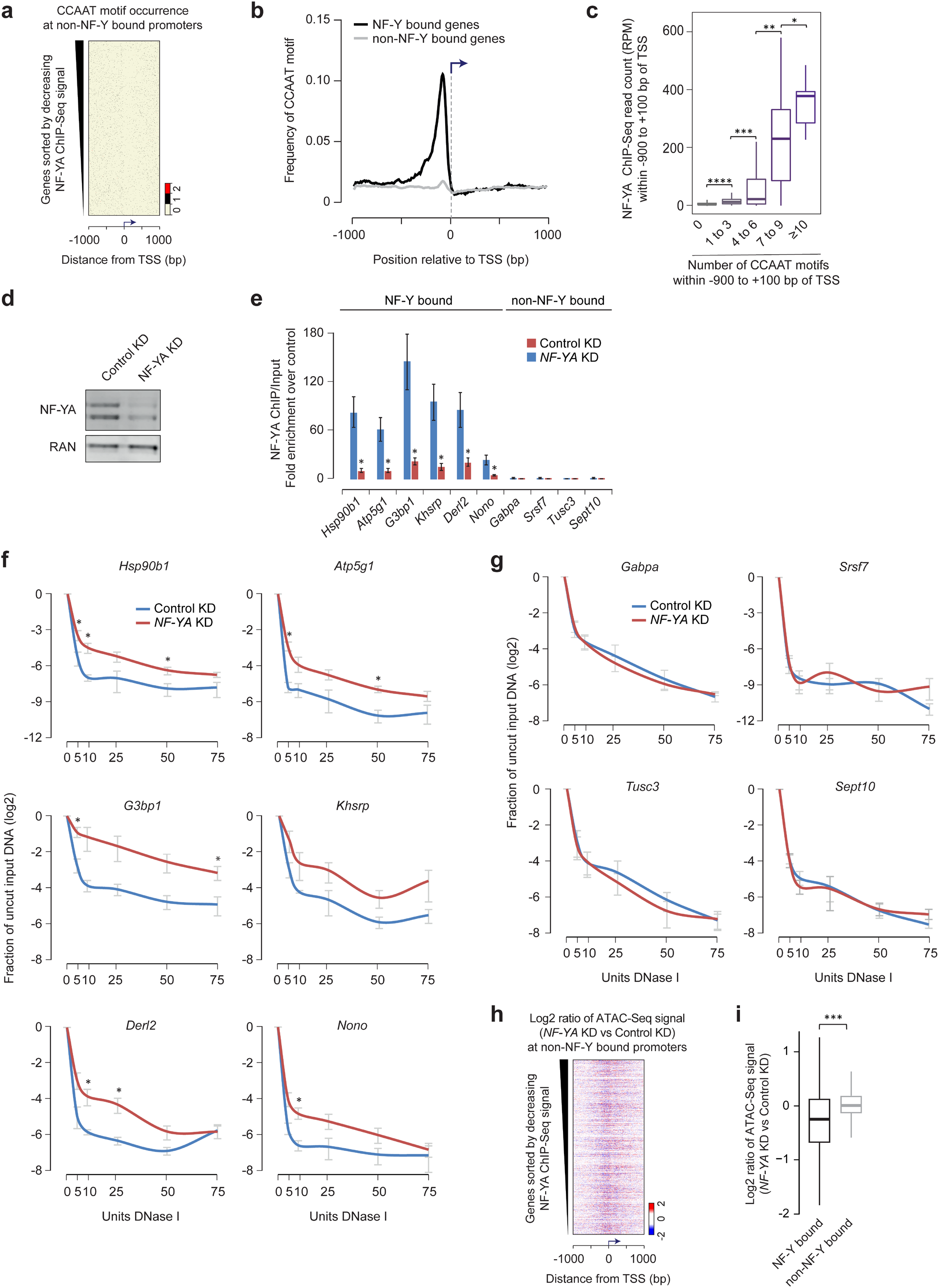
NF-Y binding is required to maintain accessible chromatin at its target promoters. **a,** CCAAT motif occurrence near TSSs of genes without promoter-proximal NF-Y binding. **b,** Frequency of CCAAT motif occurrence near TSSs of genes with (*n* = 3,056, black) or without *n* = 21,195, grey) promoter-proximal NF-Y binding in ESCs. **c,** Box plot showing the relationship between CCAAT motif occurrence within the promoter (−900 bp to +100 bp region relative to TSS, x-axis) and NF-YA ChIP-Seq read density (y-axis) for all annotated RefSeq genes. *P-value = 0.013, **P-value = 2.12E-22, ***P-value = 1.11E-88, ****P-value = 0 (Wilcoxon rank-sum test, two-sided) **d,** Western-blot analysis of NF-YA in control or *NF-YA* KD ESCs, 48 hr after siRNA transfection. Ran used as a loading control. **e,** ChIP-qPCR analysis of NF-Y binding sites in control or *NF-YA* KD ESCs. Error bars, SEM of three to five Biological replicates. **f, g** DNase I hypersensitivity and qPCR analysis of promoters with (f) or without (g) NF-Y binding in control (blue) and *NF-YA* KD (red) ESCs. Error bars, SEM of three biological replicates. *P-value < 0.05 (Student’s t-test, two-sided). **h,** Relative change (log2) in chromatin accessibility, as measured using ATAC-Seq, near TSSs of genes without promoter-proximal NF-Y binding in *NF-YA* KD vs control KD ESCs. **i,** Box plot showing the distribution of foldchanges in ATAC-Seq signal (in NF-YA KD vs control KD ESCs) within the upstream proximal-promoter regions (−150 bp to −50 bp). ***P-value = 2.86E-209 (Wilcoxon rank-sum test, two-sided)

**Supplementary Figure 2.**
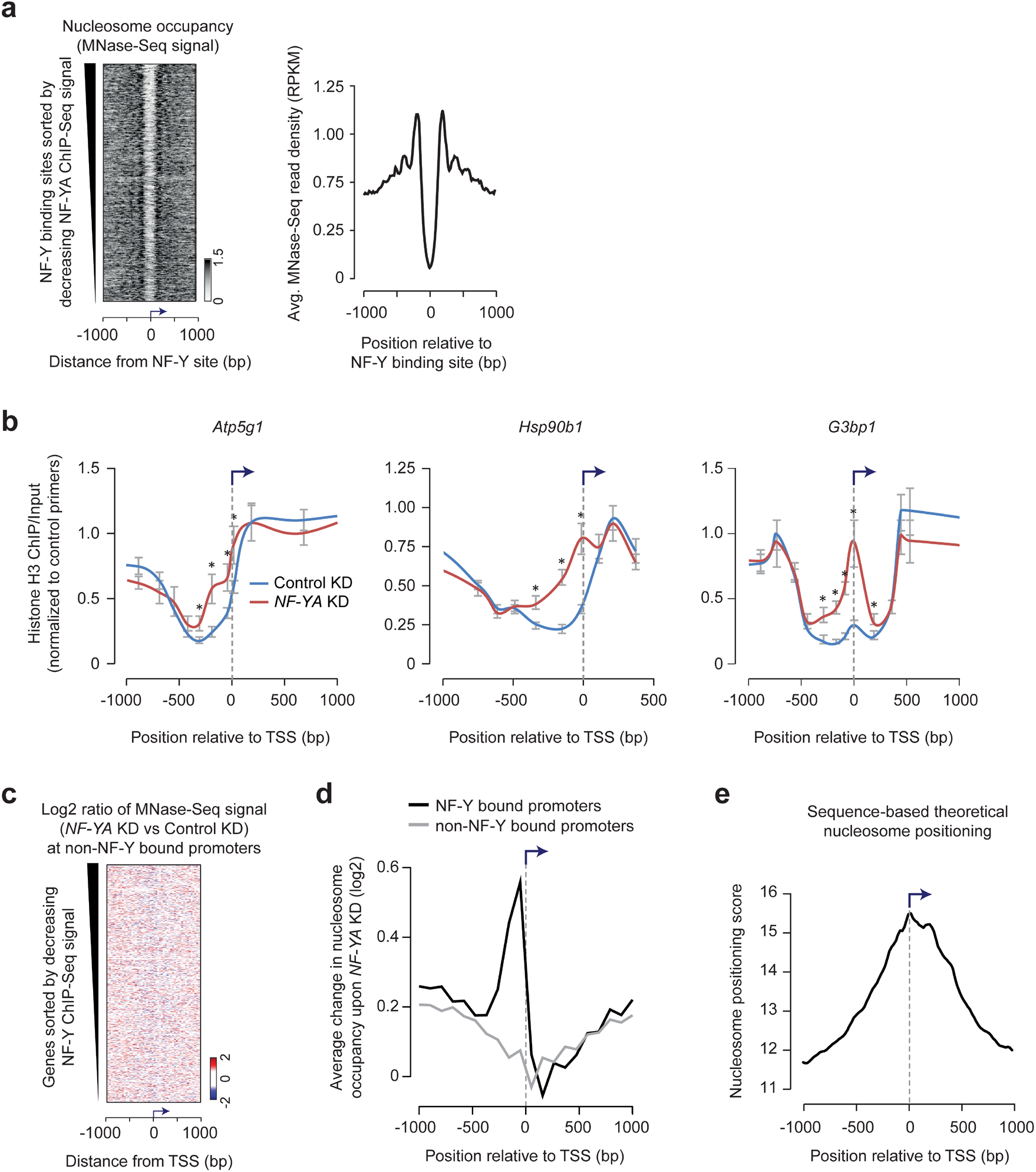
NF-Y binding protects nucleosome-depleted region from nucleosome encroachment. **a,** *Left*: Nucleosome occupancy, as measured using MNase-Seq, at all NF-Y binding sites (*n* = 5,359) in ESCs. *Right:* Average nucleosome occupancy (y-axis) at NF-Y binding sites. RPKM, reads per million mapped reads. **b,** ChIP-qPCR analysis of Histone H3 occupancy at candidate gene promoters with NF-Y binding in control or *NF-YA* KD ESCs. Error bars, SEM of three replicates. **c,** Relative change in nucleosome occupancy (gain, red; loss, blue) near TSSs of genes without promoter-proximal NF-Y binding in *NF-YA* KD vs control KD ESCs. **d,** Average change in nucleosome occupancy (y-axis) in *NF-YA* KD vs control KD ESCs near TSSs of genes with (black) or without (gray) promoter-proximal NF-Y binding. **e,** Density plot showing average theoretical nucleosome positioning, predicted based on DNA sequence composition, near TSSs of genes with promoter-proximal NF-Y binding in ESCs.

**Supplementary Figure 3.**
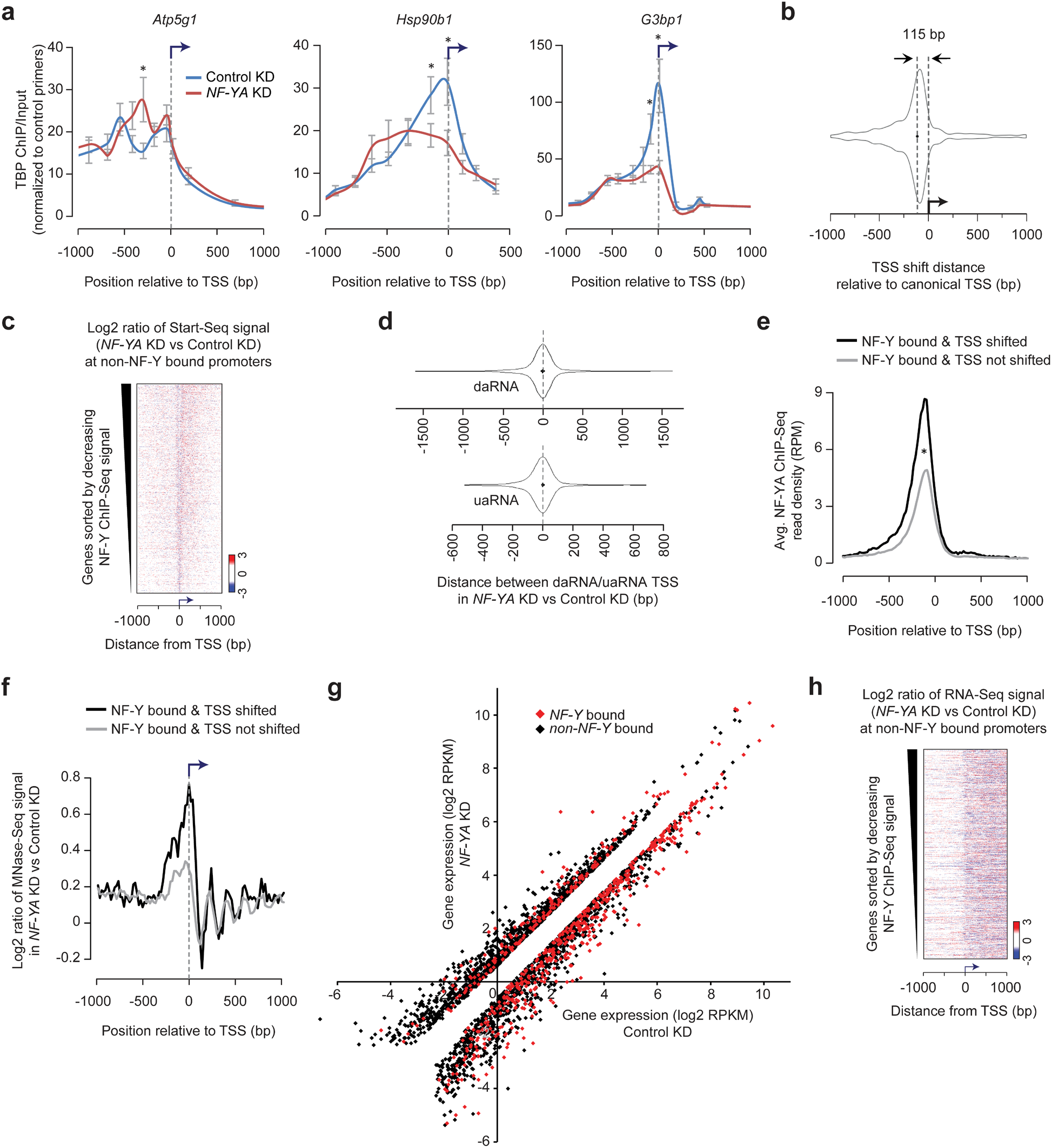
NF-Y binding influences PIC positioning and TSS selection. **a,** ChIP-qPCR analysis of TBP occupancy at candidate gene promoters with promoter-proximal NF-Y binding in control (blue) or *NF-YA* KD (red) ESCs. Error bars, SEM of three replicates. **b,** Violin plot showing the distribution of TSS shift distance, as measured using Start-Seq, in *NF-YA* KD vs control KD ESCs. Median shift distance = 115bp. **c,** Relative fold-change (log2) in Start-Seq signal (red, gain; blue, loss) in *NF-YA* KD vs control KD cells near TSSs of genes without promoter-proximal NF-Y binding. **d,** Violin plot showing the distribution of distance by which the TSS of upstream (or downstream) anti-sense RNA (uaRNA or daRNA, respectively) shifted its position, as measured using Start-Seq, in *NF-YA* KD vs control KD ESCs. Median indicated by dotted grey line. **e,** Average NF-YA occupancy (y-axis), as measured using ChIP-Seq, near TSSs of genes with promoter-proximal NF-Y binding that exhibit (black, *n* = 538) or do not exhibit (grey, *n* = 2.518) an ectopic TSS (TSS shift) in *NF-YA* KD vs control KD ESCs. RPM, reads per million mapped reads. **f,** Relative change in nucleosome occupancy (gain, red; loss, blue) near TSSs of genes with promoter-proximal NF-Y binding that exhibit (black, *n* = 538) or do not exhibit (grey, *n* = 2.518) an ectopic TSS (TSS shift) in *NF-YA* KD vs control KD ESCs. **g,** Scatter plot showing the (RNA-Seq) expression of NF-Y bound (red, *n* = 660) and non-NF-Y bound (black, *n* = 2,389) genes in control (x-axis) and NF-YA KD (y-axis) ESCs. Only genes whose expression is significantly different (>1.5-fold) in NF-YA KD vs control cells are shown. **h,** Relative fold-change (log2) in RNA-Seq signal (red, gain; blue, loss) in *NF-YA* KD vs control KD cells near TSSs of genes without promoter-proximal NF-Y binding.

**Supplementary Figure 4.**
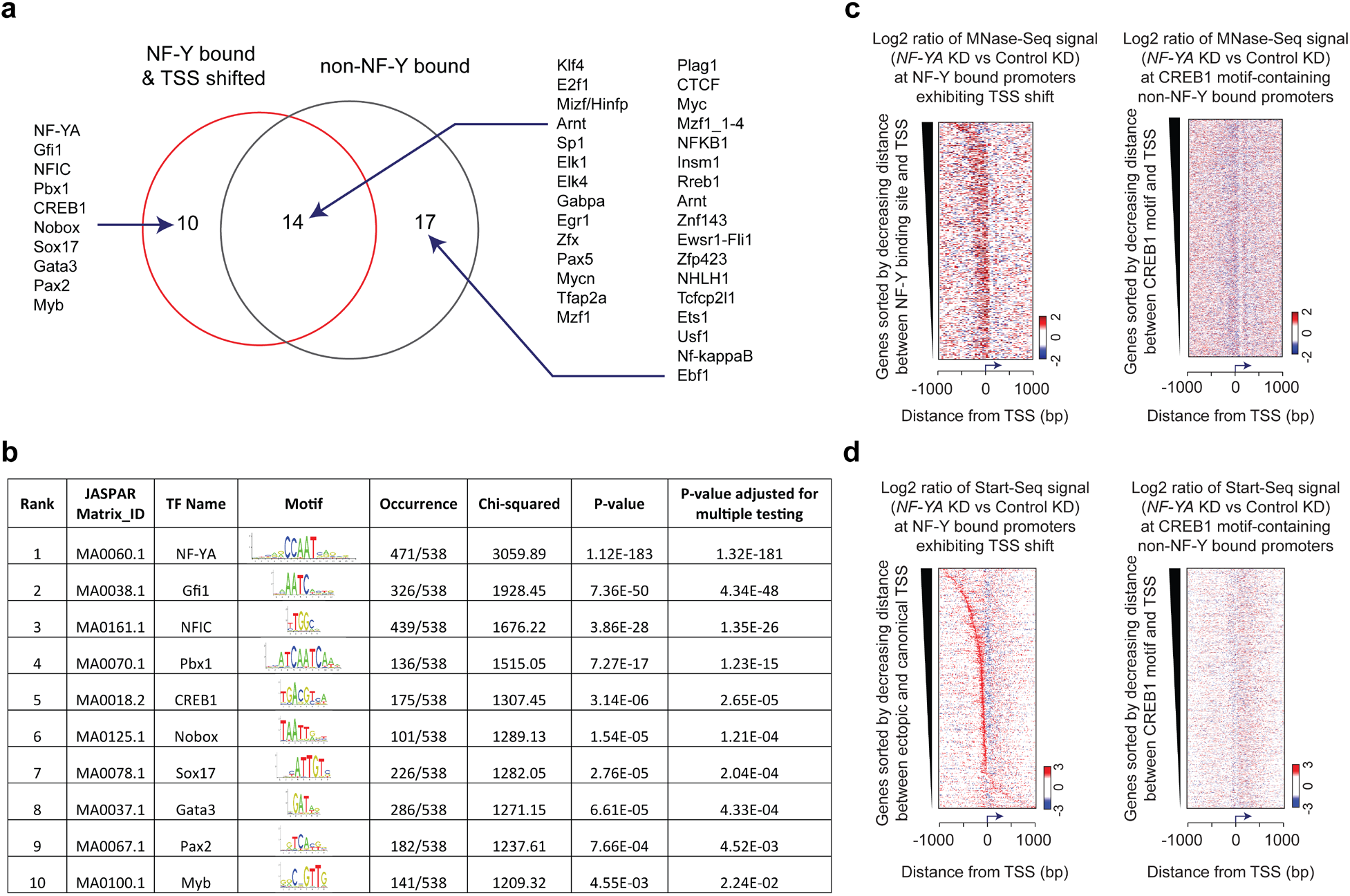
Analysis of TF binding motifs enriched within the promoters of NF-Y bound genes exhibiting a TSS shift. **a,** Venn diagram summarizing the list of TF binding motifs (source: JASPAR database) enriched within the promoter sequences (200 bp upstream of TSS) of NF-Y bound genes exhibiting a TSS shift and non-NF-Y bound genes. **b,** Summary statistics of TF binding motifs enriched within the promoter sequences (200 bp upstream of TSS) of NF-Y bound genes exhibiting a TSS shift. **c,** Comparison of changes in nucleosome occupancy (NF-YA KD vs control KD) near TSSs of NF-Y bound genes exhibiting a TSS shift (*left*) vs non-NF-Y bound genes containing a CREB1 binding motif within 200 bp upstream of TSS (*right*). **d,** Comparison of changes in Start-Seq signal (NF-YA KD vs control KD) near TSSs of NF-Y bound genes exhibiting a TSS shift (*left*) vs non-NF-Y bound genes containing a CREB1 binding motif within 200 bp upstream of TSS (*right*).

**Supplementary Figure 5.**
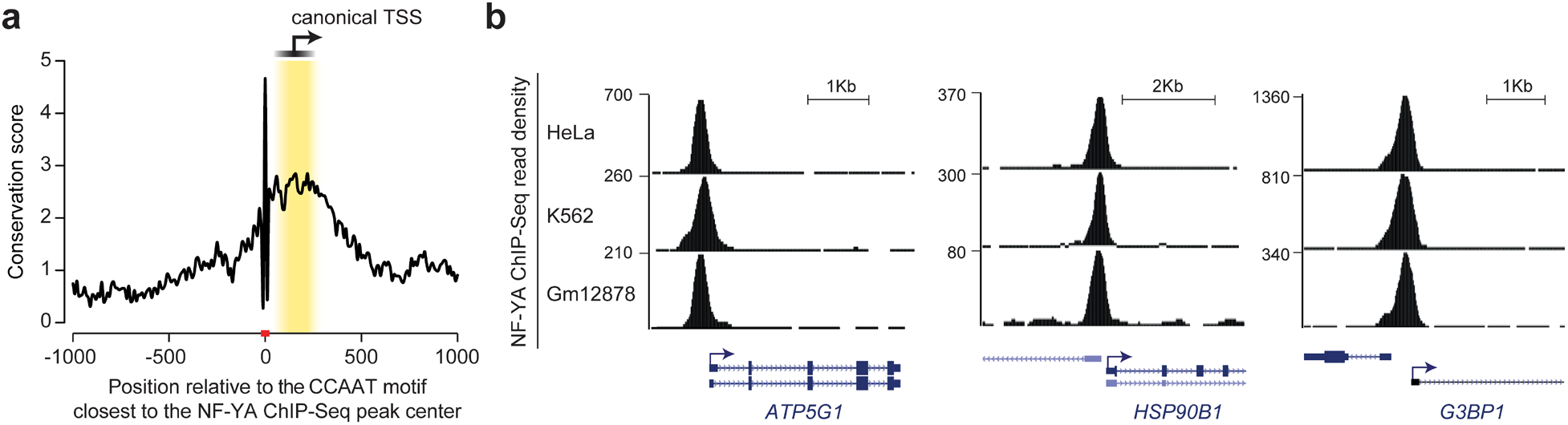
Conservation of NF-Y binding in mammals. **a,** Average sequence conservation in mammals, as computed using PhyloP, of the 2 Kb region centered on the CCAAT motif closest to the NF-Y peak center. Only genes with promoter-proximal NF-Y binding that exhibit an ectopic TSS were used (*n* = 538). **b,** Genome browser shots of genes, which bind NF-Y within promoter-proximal regions in mouse ESCs (see Figure 2), showing promoter-proximal NF-Y binding in human normal (Gm12878) and cancer cells (HeLa and K562). Human NF-YA ChIP-Seq data from ENCODE (GSE31477) was used.

**Supplementary Figure 6.**
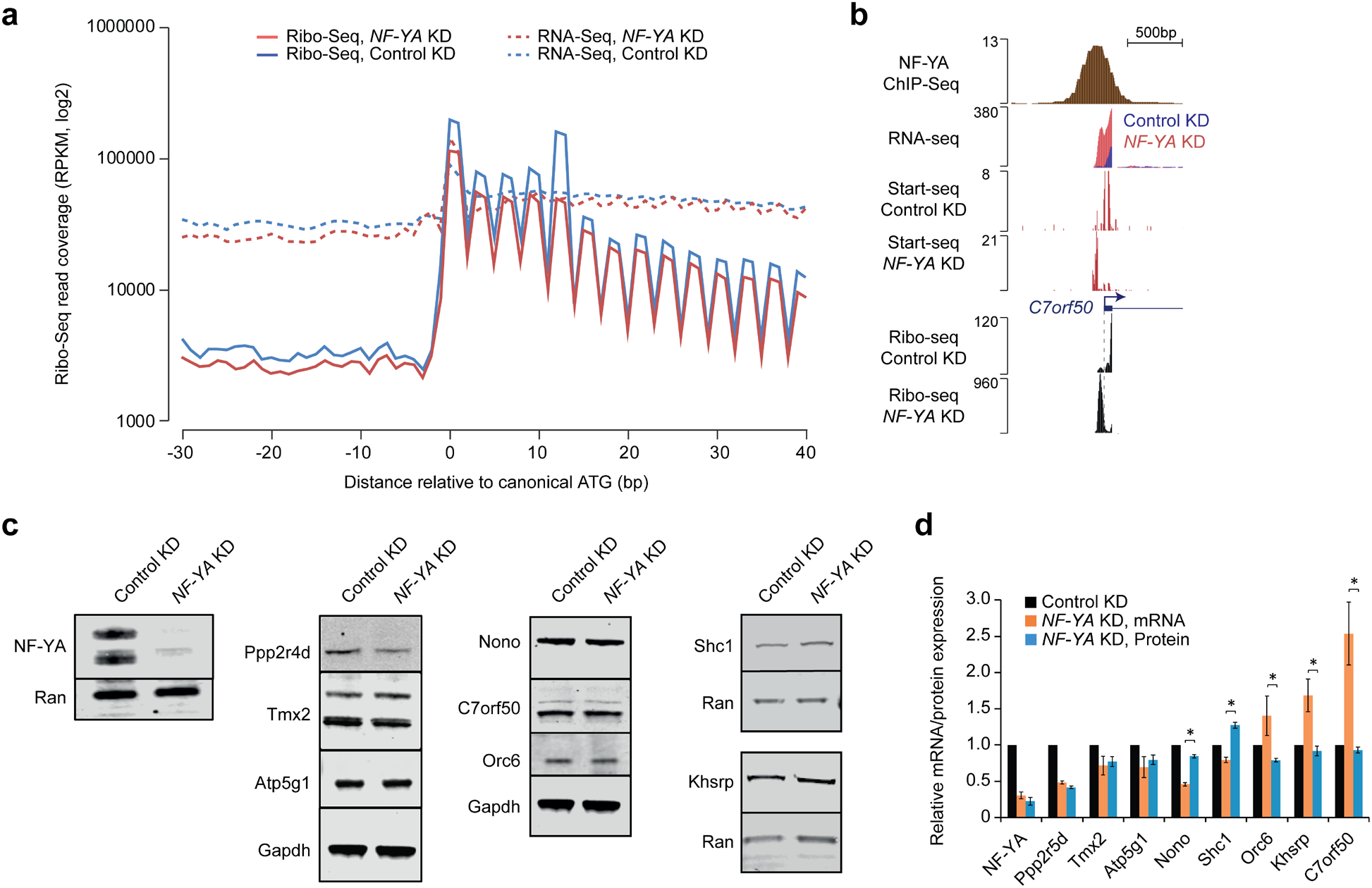
Transcripts originating from ectopic TSSs in NF-Y depleted cells undergo translation. **e,** Ribo-seq and RNA-Seq read coverage, centered on the canonical start codon (ATG), for all genes in control and *NF-YA* KD ESCs. Triplet phasing, beginning at the canonical start codon, is observed for Ribo-Seq but not RNA-Seq data. RPKM, reads per kilobase per million mapped reads. **f,** Genome browser shot of NF-Y target gene *C7orf50* (3110082I17Rik) showing ribosome-protected RNA expression, as measured using Ribo-Seq, in control (blue) and *NF-YA* KD (red) ESCs. Also shown are tracks for RNA-Seq and Start-Seq in control and *NF-YA* KD ESCs. **g,** Western-blot analysis of NF-YA, Ppp2r5d, Tmx2, Atp5g1, Nono, C7orf50, Orc6, Shc1 and Khsrp in control and NF-YA knock-down (KD) ESCs 48h after siRNA transfection. Ran or Gapdh used as loading controls. Representative images are shown. **h,** Relative changes in gene expression (mRNA, orange) and protein levels (blue) in *NF-YA* KD vs control ESCs. Protein levels were determined using Western-blot analysis (see Figure S4C), quantified by Licor Image Studio^®^ software. mRNA data normalized to *Actin, HAZ* and *TBP*. Protein data normalized to Ran or Gapdh. Error bars, SEM of three to five biological replicates. *P-value < 0.00002 (Student’s t-test, two-sided) **i,** Scatter plot showing the (RNA-Seq) expression of NF-Y bound (red) and non-NF-Y bound (black) genes in control (x-axis) and NF-YA KD (y-axis) ESCs.

## REFERENCES

1. Epstein, W. & Beckwith, J.R. Regulation of gene expression. Ann. Rev. Biochem. 37, 411–436 (1968).

2. Valen, E. & Sandelin, A. Genomic and chromatin signals underlying transcription start-site selection. Trends Genet 27, 475–85 (2011).

3. Muller, F. & Tora, L. Chromatin and DNA sequences in defining promoters for transcription initiation. Biochim Biophys Acta 1839, 118–28 (2014).

4. Core, L.J. et al. Analysis of nascent RNA identifies a unified architecture of initiation regions at mammalian promoters and enhancers. Nat Genet 46, 1311–20 (2014).

5. Roy, A.L. & Singer, D.S. Core promoters in transcription: old problem, new insights. Trends Biochem Sci 40, 165–71 (2015).

6. Vo Ngoc, L., Wang, Y.L., Kassavetis, G.A. & Kadonaga, J.T. dThe punctilious RNA polymerase II core promoter. Genes Dev 31, 1289–1301 (2017).

7. Danino, Y.M., Even, D., Ideses, D. & Juven-Gershon, T. The core promoter: At the heart of gene expression. Biochim Biophys Acta 1849, 1116–31 (2015).

8. Lai, W.K.M. & Pugh, B.F. Understanding nucleosome dynamics and their links to gene expression and DNA replication. Nat Rev Mol Cell Biol 18, 548–562 (2017).

9. Scruggs, B.S. et al. Bidirectional Transcription Arises from Two Distinct Hubs of Transcription Factor Binding and Active Chromatin. Mol Cell 58, 1101–12 (2015).

10. Smale, S.T. & Kadonaga, J.T. The RNA polymerase II core promoter. Annu Rev Biochem 72, 449–79 (2003).

11. Carninci, P. et al. Genome-wide analysis of mammalian promoter architecture and evolution. Nat Genet 38, 626–35 (2006).

12. Pinto, I., Wu, W.H., Na, J.G. & Hampsey, M. Characterization of sua7 mutations defines a domain of TFIIB involved in transcription start site selection in yeast. J Biol Chem 269, 30569–73 (1994).

13. Ghazy, M.A., Brodie, S.A., Ammerman, M.L., Ziegler, L.M. & Ponticelli, A.S. Amino acid substitutions in yeast TFIIF confer upstream shifts in transcription initiation and altered interaction with RNA polymerase II. Mol Cell Biol 24, 10975–85 (2004).

14. Kasahara, K., Ohyama, Y. & Kokubo, T. Hmo1 directs pre-initiation complex assembly to an appropriate site on its target gene promoters by masking a nucleosome-free region. Nucleic Acids Res 39, 4136–50 (2011).

15. Wang, Q. & Donze, D. Transcription factor Reb1 is required for proper transcriptional start site usage at the divergently transcribed TFC6-ESC2 locus in Saccharomyces cerevisiae. Gene (2016).

16. Venters, B.J. & Pugh, B.F. A canonical promoter organization of the transcription machinery and its regulators in the Saccharomyces genome. Genome Res 19, 360–71 (2009).

17. Rhee, H.S. & Pugh, B.F. Genome-wide structure and organization of eukaryotic preinitiation complexes. Nature 483, 295–301 (2012).

18. Maity, S.N. & de Crombrugghe, B. Role of the CCAAT-binding protein CBF/NF-Y in transcription. Trends Biochem Sci 23, 174–8 (1998).

19. Dolfini, D., Gatta, R. & Mantovani, R. NF-Y and the transcriptional activation of CCAAT promoters. Crit Rev Biochem Mol Biol 47, 29–49 (2012).

20. Maity, S.N. NF-Y (CBF) regulation in specific cell types and mouse models. Biochim Biophys Acta 1860, 598–603 (2017).

21. Nardini, M. et al. Sequence-specific transcription factor NF-Y displays histone-like DNA binding and H2B-like ubiquitination. Cell 152, 132–43 (2013).

22. Huber, E.M., Scharf, D.H., Hortschansky, P., Groll, M. & Brakhage, A.A. DNA minor groove sensing and widening by the CCAAT-binding complex. Structure 20, 1757–68 (2012).

23. Oldfield, A.J. et al. Histone-fold domain protein NF-Y promotes chromatin accessibility for cell type-specific master transcription factors. Mol Cell 55, 708–22 (2014).

24. Fleming, J.D. et al. NF-Y coassociates with FOS at promoters, enhancers, repetitive elements, and inactive chromatin regions, and is stereo-positioned with growth-controlling transcription factors. Genome Res 23, 1195–209 (2013).

25. Kaplan, N. et al. The DNA-encoded nucleosome organization of a eukaryotic genome. Nature 458, 362–6 (2009).

26. Imbalzano, A.N., Kwon, H., Green, M.R. & Kingston, R.E. Facilitated binding of TATA-binding protein to nucleosomal DNA. Nature 370, 481–5 (1994).

27. Bellorini, M. et al. CCAAT binding NF-Y-TBP interactions: NF-YB and NF-YC require short domains adjacent to their histone fold motifs for association with TBP basic residues. Nucleic Acids Res 25, 2174–81 (1997).

28. Core, L.J., Waterfall, J.J. & Lis, J.T. Nascent RNA sequencing reveals widespread pausing and divergent initiation at human promoters. Science 322, 1845–8 (2008).

29. Seila, A.C. et al. Divergent transcription from active promoters. Science 322, 1849–51 (2008).

30. Preker, P. et al. RNA exosome depletion reveals transcription upstream of active human promoters. Science 322, 1851–4 (2008).

31. Siepel, A. et al. Evolutionarily conserved elements in vertebrate, insect, worm, and yeast genomes. Genome Res 15, 1034–50 (2005).

32. Dikstein, R. Transcription and translation in a package deal: the TISU paradigm. Gene 491, 1–4 (2012).

33. Ingolia, N.T., Ghaemmaghami, S., Newman, J.R. & Weissman, J.S. Genome-wide analysis in vivo of translation with nucleotide resolution using ribosome profiling. Science 324, 218–23 (2009).

34. Jovanovic, M. et al. Immunogenetics. Dynamic profiling of the protein life cycle in response to pathogens. Science 347, 1259038 (2015).

35. Liu, Y. et al. Impact of Alternative Splicing on the Human Proteome. Cell Rep 20, 1229–1241 (2017).

36. Wethmar, K. The regulatory potential of upstream open reading frames in eukaryotic gene expression. Wiley Interdiscip Rev RNA 5, 765–78 (2014).

37. Johnstone, T.G., Bazzini, A.A. & Giraldez, A.J. Upstream ORFs are prevalent translational repressors in vertebrates. EMBO J 35, 706–23 (2016).

38. Cheng, Z. et al. Pervasive, Coordinated Protein-Level Changes Driven by Transcript Isoform Switching during Meiosis. Cell 172, 910–923 e16 (2018).

39. Chen, J. et al. Kinetochore inactivation by expression of a repressive mRNA. Elife 6(2017).

40. Chia, M. et al. Transcription of a 5’ extended mRNA isoform directs dynamic chromatin changes and interference of a downstream promoter. Elife 6(2017).

41. Sherwood, R.I. et al. Discovery of directional and nondirectional pioneer transcription factors by modeling DNase profile magnitude and shape. Nat Biotechnol 32, 171–8 (2014).

42. Lu, F. et al. Establishing Chromatin Regulatory Landscape during Mouse Preimplantation Development. Cell 165, 1375–88 (2016).

43. Tao, Z. et al. Embryonic epigenetic reprogramming by a pioneer transcription factor in plants. Nature 551, 124–128 (2017).

44. Romier, C., Cocchiarella, F., Mantovani, R. & Moras, D. The NF-YB/NF-YC structure gives insight into DNA binding and transcription regulation by CCAAT factor NF-Y. J Biol Chem 278, 1336–45 (2003).

45. Liberati, C., di Silvio, A., Ottolenghi, S. & Mantovani, R. NF-Y binding to twin CCAAT boxes: role of Q-rich domains and histone fold helices. J Mol Biol 285, 1441–55 (1999).

46. Coustry, F., Hu, Q., de Crombrugghe, B. & Maity, S.N. CBF/NF-Y functions both in nucleosomal disruption and transcription activation of the chromatin-assembled topoisomerase IIalpha promoter. Transcription activation by CBF/NF-Y in chromatin is dependent on the promoter structure. J Biol Chem 276, 40621–30 (2001).

47. Motta, M.C., Caretti, G., Badaracco, G.F. & Mantovani, R. Interactions of the CCAAT-binding trimer NF-Y with nucleosomes. J Biol Chem 274, 1326–33 (1999).

48. Kukimoto, I., Elderkin, S., Grimaldi, M., Oelgeschlager, T. & Varga-Weisz, P.D. The histone-fold protein complex CHRAC-15/17 enhances nucleosome sliding and assembly mediated by ACF. Mol Cell 13, 265–77 (2004).

49. Frontini, M. et al. NF-Y recruitment of TFIID, multiple interactions with histone fold TAF(II)s. J Biol Chem 277, 5841–8 (2002).

50. Dorn, A. et al. Conserved major histocompatibility complex class II boxes--X and Y--are transcriptional control elements and specifically bind nuclear proteins. Proc Natl Acad Sci U S A 84, 6249–53 (1987).

51. Rach, E.A. et al. Transcription initiation patterns indicate divergent strategies for gene regulation at the chromatin level. PLoS Genet 7, e1001274 (2011).

52. Asada, R., Takemata, N., Hoffman, C.S., Ohta, K. & Hirota, K. Antagonistic controls of chromatin and mRNA start site selection by Tup family corepressors and the CCAAT-binding factor. Mol Cell Biol 35, 847–55 (2015).

53. Segal, E. et al. A genomic code for nucleosome positioning. Nature 442, 772–8 (2006).

54. Lobanov, A.V., Turanov, A.A., Hatfield, D.L. & Gladyshev, V.N. Dual functions of codons in the genetic code. Crit Rev Biochem Mol Biol 45, 257–65 (2010).

55. Starck, S.R. et al. Leucine-tRNA initiates at CUG start codons for protein synthesis and presentation by MHC class I. Science 336, 1719–23 (2012).

56. Dever, T.E. Molecular biology. A new start for protein synthesis. Science 336, 1645–6 (2012).

57. Calvo, S.E., Pagliarini, D.J. & Mootha, V.K. Upstream open reading frames cause widespread reduction of protein expression and are polymorphic among humans. Proc Natl Acad Sci U S A 106, 7507–12 (2009).

58. Vilela, C. & McCarthy, J.E. Regulation of fungal gene expression via short open reading frames in the mRNA 5’ untranslated region. Mol Microbiol 49, 859–67 (2003).

59. Cinghu, S. et al. Intragenic Enhancers Attenuate Host Gene Expression. Mol Cell 68, 104–117 e6 (2017).

60. Williams, L.H. et al. Pausing of RNA polymerase II regulates mammalian developmental potential through control of signaling networks. Mol Cell 58, 311–322 (2015).

61. Langmead, B., Trapnell, C., Pop, M. & Salzberg, S.L. Ultrafast and memory-efficient alignment of short DNA sequences to the human genome. Genome Biol 10, R25 (2009).

62. Kim, D. et al. TopHat2: accurate alignment of transcriptomes in the presence of insertions, deletions and gene fusions. Genome Biol 14, R36 (2013).

63. Trapnell, C. et al. Differential gene and transcript expression analysis of RNA-seq experiments with TopHat and Cufflinks. Nat Protoc 7, 562–78 (2012).

64. Love, M.I., Huber, W. & Anders, S. Moderated estimation of fold change and dispersion for RNA-seq data with DESeq2. Genome Biol 15, 550 (2014).

65. Patro, R., Duggal, G., Love, M.I., Irizarry, R.A. & Kingsford, C. Salmon provides accurate, fast, and bias-aware transcript expression estimates using dual-phase inference. bioRxiv (2016).

66. Nechaev, S. et al. Global analysis of short RNAs reveals widespread promoter-proximal stalling and arrest of Pol II in Drosophila. Science 327, 335–8 (2010).

67. Henriques, T. et al. Stable pausing by RNA polymerase II provides an opportunity to target and integrate regulatory signals. Mol Cell 52, 517–28 (2013).

68. Martin, M. Cutadapt removes adapter sequences from high-throughput sequencing reads. 2011 17(2011).

69. Anders, S. & Huber, W. Differential expression analysis for sequence count data. Genome Biol 11, R106 (2010).

70. Dobin, A. et al. STAR: ultrafast universal RNA-seq aligner. Bioinformatics 29, 15–21 (2013).

71. Bailey, T.L. et al. MEME SUITE: tools for motif discovery and searching. Nucleic Acids Res 37, W202–8 (2009).

72. Thomas-Chollier, M. et al. Transcription factor binding predictions using TRAP for the analysis of ChIP-seq data and regulatory SNPs. Nat Protoc 6, 1860–9 (2011).

73. Mathelier, A. et al. JASPAR 2016: a major expansion and update of the open-access database of transcription factor binding profiles. Nucleic Acids Res 44, D110–5 (2016).

74. Pollard, K.S., Hubisz, M.J., Rosenbloom, K.R. & Siepel, A. Detection of nonneutral substitution rates on mammalian phylogenies. Genome Res 20, 110–21 (2010).

